# EpiModel: An R Package for Mathematical Modeling of Infectious Disease over Networks

**DOI:** 10.1101/213009

**Authors:** Samuel M. Jenness, Steven M. Goodreau, Martina Morris

**Affiliations:** Emory University, Department of Epidemiology, 1518 Clifton Road, Atlanta, GA 30322;; URL: http://samueljenness.org/; University of Washington

**Keywords:** mathematical model, infectious disease, epidemiology, networks, R.

## Abstract

**EpiModel** provides tools for building, simulating, and analyzing mathematical models for the population dynamics of infectious disease transmission in R. Several classes of models are included, but the unique contribution of this software package is a general stochastic framework for modeling the spread of epidemics on networks. This framework integrates recent advances in statistical methods for network analysis, temporal exponential random graph models, which allows the epidemic modeling to be firmly grounded in empirical data on the contacts and persistent partnerships that can spread infection. This article provides an overview of both the modeling tools built into **EpiModel**, designed to facilitate learning for students new to modeling, and the application programming interface for extending **EpiModel**, designed to facilitate the exploration of novel research questions for advanced modelers.

## 1. Introduction

The **EpiModel** package provides tools for building, simulating, and analyzing mathematical models for epidemics using R. Epidemic models are a formal representation of the three basic processes that collectively determine the population dynamics of infectious disease transmission: the contact process, the infection process (between- and within-host), and the demographic process. The *structure* of the model determines how these processes interact at the micro-level to produce the macro-level outcomes of disease incidence and prevalence (Anderson and May 1992). The *input parameters* determine the rates or probabilities of events occurring in these processes; they can be estimated from data, treated as sensitivity ranges to examine the impact on macro-level model outcomes, or used for calibrating model outcomes to observed data. In contrast to traditional statistical models (e.g., the **surveillance** (Höhle *et al*. 2016), **epifit** (Doi *et al.* 2016) and **EpiEstim** (Cori 2013) packages in R), which estimate parameters from empirical data, mathematical epidemic models use already estimated parameters to produce simulated data (e.g., the **epinet** (Groendyke and Welch 2016) and **EpiDynamics** (Santos Baquero and Silveira Marques 2015) packages). This provides an *in silico* laboratory to gain insight into the dynamics of these complex systems, test empirical hypotheses about the determinants of a specific outbreak trajectory, and forecast the impact of interventions like vaccines, clinical treatment, or public health education campaigns (Garnett *et al*. 2011; Lessler and Cummings 2016).

A range of different modeling tools has been developed in the field of mathematical epidemiology over the last century. All modeling approaches represent the population stratified by state of infection (susceptible and infected, at a minimum) and a process of infection (the transition between these states). Beyond this there is tremendous variation in the heterogeneity in states and processes represented, the mathematical theory and methods used, the integration of the statistical and mathematical modeling, and the strengths and limitations that follow.

**EpiModel** includes support to learn, build, and simulate three distinct classes of epidemic models:

1. *Deterministic compartmental models (DCMs)* are based on systems of differential equations for the movement of the population through discrete states, including entry into and exit from the population, at specified rates. DCMs are the most commonly used model class in the field of mathematical epidemiology today, in part because simple formulations can be solved analytically, or with minimal computational burden. They can represent discrete forms of heterogeneity in the population, and have a limited ability to represent persistent partnerships. With DCMs, once the structure and parameters have been specified, there is no variation in model outcomes.
2. *Stochastic individual contact models (ICMs)*, also known as *individual-based* or *agent-based* models, explicitly represent individual units in the population and the contacts between them as unique, discrete events. Compared to DCMs, they allow for more heterogeneity in specifying the contact process and other epidemiologically relevant events, and their stochasticity provides information on the range of plausible outcomes resulting from a given set of parameters. Drawbacks of these models include the potentially large amounts of input data needed for parameterization and the computational burden associated with running multiple stochastic simulations.
3. *Network models* are also stochastic and represent individual units, but unlike ICMs, they provide a general, flexible framework for representing repeated contacts with the same person or persons over time (e.g., sexual partnerships). These repeated contacts give rise to persistent network configurations – pairs, triples, and larger connected components – that in turn may establish the temporally ordered pathways for infectious disease transmission across a population. **EpiModel** uses recently developed statistical methods for network analysis to provide a generalized framework for both estimation and simulation of dynamically evolving networks with specifiable configurations, and the acts within partnerships that transmit infection. Network models provide the most accurate control over the contact process, but have greater computational burden than ICMs, both because they require statistical estimation of the network model parameters (for a true network model) and because the simulation algorithm must address a complex set of overlapping constraints.

**EpiModel** provides functions for each of these model classes, but the network models are the unique contribution of this software package. Interest in network models has grown rapidly, largely driven by the global heterogeneity observed in the epidemics of human immunodeficiency virus (HIV) and other sexually transmitted infections (STIs). Both the transmission dynamics and the prevalence of disparities in HIV are highly sensitive to variations in the underlying partnership network structure, as shown by Morris *et al*. (2009) for HIV/STIs in the United States. Recent developments in statistical methods for network analysis now provide a principled approach to estimating and simulating this partnership network structure for epidemic modeling purposes, which require dynamic networks where not only the edges (partnerships) change but also the number and attributes of the nodes (persons) (Krivitsky *et al.* 2011; Krivitsky and Handcock 2014).

Representing the details of the *repeated* contact structure – the timing and sequence of partnerships, and the acts within them – is most important when transmission requires intimate contact, the intimate contact is relatively rare, and the probability of infection per contact is relatively low (Goodreau 2011). In this context, the network connectivity needed for epidemic persistence may not be captured by summaries like the rate of partner acquisition over time (Carnegie and Morris 2012). Instead, the temporal overlaps in partnerships (“concurrency”) may provide the connectivity needed to sustain transmission over time (Morris and Kretzschmar 2000).

DCMs were not originally designed to represent these types of partnership networks. They may be useful for investigating other components relevant for HIV/STI contact processes, including heterogeneity in contact rates (and mixing between high-risk and low-risk groups), which was shown in early modeling studies to enable endemic persistence of STIs that would not be predicted by the simple homogenous model (Hethcote and Van Ark 1987). However, this type of “core group” heterogeneity does not appear to explain the generalized epidemics of HIV, as the behaviors observed in these populations are profoundly different than the behavioral model specifications required for simulations to replicate observed disease prevalence (Garnett and Anderson 1993; Hallett *et al*. 2008; Abu-Raddad and Longini 2008).

Network models, in contrast, have been able to replicate generalized HIV epidemic dynamics when driven with observed behavioral data and realistic transmission parameters (Jenness *et al.* 2016b). The partnership networks produced by these models emerge from an empirically informed set of micro-level behaviors, including (but not limited to) how people choose their partners (e.g., based on attributes like sex and age), how many partners persons have at any one time, and the distribution of partnership lengths and overlaps (Goodreau *et al*. 2012; Jenness *et al*. 2016a). These behaviors can be measured through survey data collection, statistically analyzed in a way that jointly estimates the multiple correlated underlying parameters, and the statistical model can then be used to drive a network-based epidemic simulation (Morris 1997; Goodreau *et al*. 2010). The statistical theory that makes this possible was developed in the literature on exponential-family random graph models (ERGMs) (Holland and Leinhardt 1981; Frank and Strauss 1986; Hunter and Handcock 2006) and the recent extensions to temporal ERGMs (TERGMs) (Krivitsky and Handcock 2014). This kind of principled, integrated network estimation/simulation framework is new in epidemic modeling, and it is the basis of the **EpiModel** software.

Both the estimation and simulation of the dynamic network in EpiModel are implemented using the Markov Chain Monte Carlo (MCMC) algorithm functions from the statnet package (Hunter *et al*. 2008). The MCMC algorithm exploits a key property of exponential family models, that the maximum likelihood estimates of the model parameters uniquely reproduce the model statistics in expectation. This is also what allows the fitted model to be used for the mathematical simulation of the contact network over time – it is theoretically guaranteed to vary stochastically around the observed network statistics (as long as the model is not degenerate). Other approaches to modeling epidemics on networks have been developed (Leung *et al*. 2015; Keeling and Rohani 2008), including applications that have used static ERGMs with a separately derived and estimated process for edge dynamics (Khan *et al.* 2014). However, TERGMs provide the only integrated, principled framework for the estimation of network models from sampled data (Krivitsky and Morris 2017) and simulation of complex dynamic networks with theoretically justified methods for handling changes in population size and composition over time (Krivitsky *et al.* 2011).

**EpiModel** has been designed to be used for both teaching purposes and advanced scientific research. The package includes a set of built-in, or *base*, model types for each of the three epidemic modeling classes, intended primarily for pedagogical purposes – these provide a comprehensive set of tools for learning alternative frameworks for basic epidemic modeling. **EpiModel** also provides an application programming interface (API) for each model class that allows users to develop modular extensions to the base models. This API was designed to facilitate scientific research.

This paper introduces the base models and the API for extensions in **EpiModel**, focusing on the stochastic network modeling class. The overall organization of the software, including functionality and tools for the other modeling classes, is described briefly in the Section 2, but the remaining sections focus on the stochastic network modeling class.

This **EpiModel** tutorial is divided into five further sections:

**Section 2** outlines the methods for accessing and getting oriented to the software, including help documentation, and provides an overview of the complete **EpiModel** package.

**Section 3** provides the theoretical and mathematical framework for modeling dynamic networks using the statistical framework of temporal ERGMs.

**Section 4** demonstrates the *base* epidemic model types built into the software. Base models provide some choice of *disease states*, and require specification of parameters in existing *modules* – self-contained functions that perform one aspect of the epidemic simulation. Two stochastic network models examples are presented to demonstrate the core functionality and options.

**Section 5** outlines the generalized **EpiModel** API for extending the disease states and modules to address new research questions. Extension modules can be written to either modify the existing functions or to build entirely new processes into the epidemic system. We show examples of both.

**Section 6** puts these tools into a broader context by discussing some of the ongoing research applications, methodological and computational challenges in this work, and opportunities for future developments.

## 2. Orientation

### 2.1. Code base

**EpiModel** is an open-source software package for the `R` computing platform. It depends on and is an extension of the **statnet** (Handcock *et al.* 2015) suite of packages for `R` developed for the representation, statistical modeling, analysis, and visualization of network data (Handcock *et al.* 2008). Our software website for **EpiModel** (http://epimodel.org/) provides a wealth of supporting documentation, tutorials, and teaching materials for the project.

Because the package is developed in an open-source framework, users may view the source code on our Github repository (http://github.com/statnet/EpiModel). Through the repository, users may submit issues for feature requests and bug reports. Advanced users may contribute to the code base by forking the public repository and submitting pull requests for changes to the code using standard Github methods.

### 2.2. Setup

To start the software demonstration, we install and load the **EpiModel** package in R, along with all dependencies. This paper was written with the current software release, **EpiModel 1.5.0**, which is hosted on CRAN, the central publication hub for `R` packages:

~~~
*R> install.packages("EpiModel", dependencies = TRUE*)
*R> library("EpiModel"*)
~~~

The main help documentation is available with the standard help methods in R. An orienting help file for the entire package may be viewed with `help("EpiModel")`. The supported model classes are briefly described along with the built-in epidemic model types across models. The overall listing of help documentation may be retrieved as follows.

~~~
*R> help(package = "EpiModel"*)
~~~

## 2.3. Software organization

Figure 1 provides an overview of the design of **EpiModel** and highlights the components that will be presented in this paper (shown with dashed box borders). As described in Section 1, **EpiModel** includes three *classes* of models but only network models will be covered since we have implemented the other two classes primarily as a gateway to learning network models. Extensive tutorials and documentation for the other classes are provided within the package and on our software website (http://epimodel.org/). This includes workshop notes for a week-long epidemic modeling course, Network Modeling for Epidemics.

**Figure 1:**
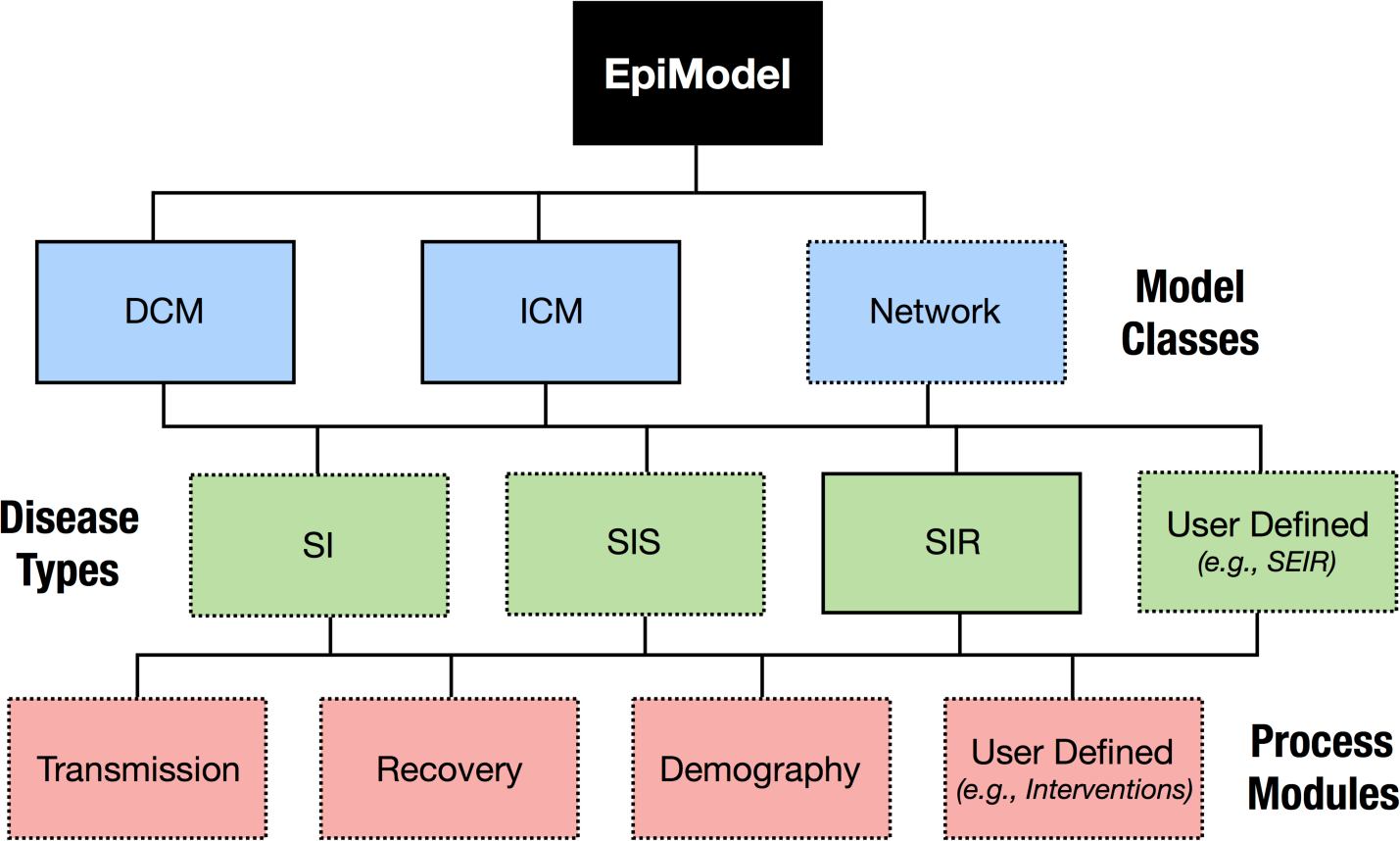
Tree diagram of **EpiModel** design. Dashed boxes are topics covered in this paper.

Within each of the classes, **EpiModel** includes three pre-structured model types, called *base* models, that facilitate the simulation of epidemic models. The primary aim of these built-in models is to reduce the programming burden for users new to epidemic modeling. These users only need to select the right model type, and learn the relevant epidemic parameters governing the processes for that model, such as rates of infection transmission or disease recovery. Users also control the population size, number of time steps for the simulation, and the number of simulations to run. It is not necessary, however, to specify the algorithms that drive the components of the system. The three types are:

- **Susceptible-Infectious (SI):** A two-state model with a one-way transition from susceptible to infected, and no recovery after infection. HIV or herpes simplex virus type 2 (HSV-2) are examples.
- **Susceptible-Infectious-Susceptible (SIS):** A two-state model in which recovery does not induce immunity, so individuals may transition back and forth between the susceptible and infected states. Examples include the common cold and curable STIs like gonorrhea.
- **Susceptible-Infectious-Recovered/Immune (SIR):** A three-state model with an additional one-way transition from infected to recovered with immunity. A classic example is measles.

Examples within this paper will highlight the SI and SIS types, while the package documentation includes additional examples for the SIR type. Experimenting with base models provides a natural learning path for developing the skills needed to program model extensions for research purposes (discussed in Section 5). These extensions involve modifying existing *process modules* that perform one component of the simulation system (disease transmission, disease progression or recovery, or demographic dynamics like births or migration). Users may also add entirely new disease types or modules that perform features not part of the base models: for example, adding stages of infection to represent variable transmissibility or waning immunity, behavioral or biological heterogeneity in transmission risk, and public health or clinical interventions. This paper will provide examples of changing the existing modules in basic ways, and Section 6 will provide references for detailed HIV/STI models with module templates that may be used for those diseases.

*Shiny Web Applications*

**EpiModel** also includes three web-based applications using the Chang *et al*. (2016) platform in R. These are primarily intended for teaching purposes, to introduce epidemic models from each class with no coding required. The three applications may be run after loading **EpiModel** through:

~~~
*R> epiweb("dcm"*)
*R> epiweb("icm"*)
*R> epiweb("net"*)
~~~

While the apps provide limited access to the range of **EpiModel** utilities, they may still be sufficient for undergraduate teaching purposes, a short overview of modeling in a larger epidemiology course, or a presentation of an epidemic simulation in real time. The rest of the tutorial presented here will focus on **EpiModel**’s command-line interface.

## 3. Network modeling framework

Network models explicitly represent discrete persons (nodes) and partnerships (edges), which are pairs of persons who remain in contact over some interval of time. Partnerships have duration, which allows for repeated acts with the same person, specification of partnership formation and dissolution rates, and control over the temporal sequencing of multiple partnerships.

This is handled through specification of a stochastic network model that captures the generative micro-level processes governing edge formation and dissolution. The approach is sometimes referred to as *bottom-up* modeling: the focus is on representing the micro-level processes (e.g., preferences for monogamy and partners from a specific age group) that lead to emergent properties at the macro-level (e.g., the global network component sizes and geodesic distributions). These macro-level network properties may be the key features that drive the dynamics of disease transmission, so they are of considerable interest.

### 3.1. Microsimulation features

To introduce the concepts and terminology for simulating network models of epidemics, it is necessary to describe three key features of the microsimulation framework for both base models and extensions. This will be especially relevant for readers with a background in DCM modeling:

1. **Units are individuals.** These models simulate epidemic spread over a population of discrete, individually identifiable elements; in contrast, DCMs treat the population as a continuous mass which is infinitely divisible, with individual elements neither identifiable nor accessible for modeling. The implications of this may be relatively minor if the stochastic simulations occur on a sufficiently large population, but there are some critical differences shown below, including the representation of continuous attributes of persons and multiple persistent partnerships.
2. **Rates and risks are stochastic.** The individual transitions between states are parameterized as random draws from distributions summarized by key parameters. In some cases, the distribution takes a simple common form like Bernoulli, binomial, uniform, or Poisson. In other cases, like the partnership formation and dissolution processes, the distribution is more complex.
3. **Time is discrete.** These models are represented in discrete time, in contrast to the continuous time algorithms typically used to solve DCMs written with differential equations. In a discrete time model, many different events can happen in one time step, and they happen in a sequence. There are no instantaneous, independent events as in continuous time. If the time step is too large relative to the rates of change, bias may be introduced into the models. Choice of time step in network models is typically a trade-off between reducing this bias with smaller steps and increased computational efficiency with larger steps.

### 3.2. Exponential random graph models

**EpiModel** depends on **statnet**, a suite of `R` packages (Handcock *et al*. 2015) that provide a robust software implementation for both static and temporal ERGMs, with a broad range of descriptive and statistical analysis tools, model fitting diagnostics, graphical utilities for network visualization, and methods for network simulation from fitted or theoretical models (Handcock *et al*. 2008). This section provides a brief review of the statistical theory implemented in **statnet** and how it is integrated into **EpiModel** through the examples below. This paper does not provide a comprehensive review of this literature or a tutorial on the packages within **statnet**. For that, the full special issue in this journal devoted to these tools (Handcock *et al.* 2008) is a good place to start, and additional details are provided in the references cited below.

**statnet** implements ERGMs to estimate and simulate relational networks based on observed patterns of density, degree, triad closure, assortivity, and other network features (Wasserman and Pattison 1996; Hunter *et al*. 2008). These methods were originally developed for modeling spatial dependence (Besag 1974), and have evolved to become the primary method for modeling dependence in networks (Holland and Leinhardt 1981; Fienberg and Wasserman 1981; Frank and Strauss 1986; Hunter and Handcock 2006). Models are typically estimated using computationally intensive MCMC algorithms (Geyer and Thompson 1992).

When using ERGMs, the edges relevant for infectious disease transmission are, in network terminology, binary and undirected. For this type of edge, ERGMs represent the probability distribution of the network **Y** as an exponential function of a set of network-level model statistics:

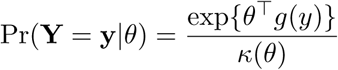

where **y** is the observed network (composed of edges, nodes, and nodal attributes), *θ* is a vector of model coefficients, *g*(*y*) is a vector of network statistics, and *κ*(*θ*) = Σ_*y′ ∈Y*_ exp{*θg*(*y′*)} is a normalizing constant representing the set of all possible configurations of a network with the size and nodal composition of **y**.

The network statistics that can be specified on the right hand side range from a single term to capture the density of the network (e.g., a Bernoulli random graph model) to a term for each dyad in the network. Each term in a model represents a “configuration” – a specific node-edge structure – and at the network level the coefficient represents how much more or less prevalent this structure is than expected (given the other terms in the model). Parsimonious models typically include a mix of *dyad-independent* and *dyad-dependent* terms. The distinction is based on whether the status of one dyad influences the status of others. Dyad dependence should be modeled with care, as the patterns it induces can cascade through the network in unexpected ways, and poorly specified models can often lead to an outcome known as “model degeneracy” (Handcock *et al*. 2003; Schweinberger 2011). With networks of any nontrivial size, *κ*(*θ*) may not be calculated analytically, and therefore simulation-based estimation procedures (MCMC) are used.

ERGMs may be reexpressed in a conditional logit form that allows for a micro-level interpretation of the parameters:

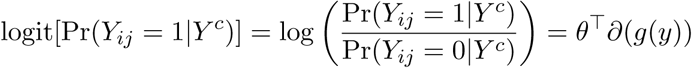

where *Y^c^* is the rest of the network, excluding *Y_ij_*, and *∂*(*g*(*y*)) are *change statistics* that represent how the count of configurations changes when *Y_ij_* is toggled from 0 to 1. The interpretation of *θ* in this expression is clearer, as it represents the conditional log-odds of an edge as a function of the number of configurations it will create.

Since epidemic models are dynamic, evolving systems, we use temporal ERGMs (TERGMs) to model the underlying dynamic network. Specifically we use separable TERGMs (Krivitsky and Handcock 2014), in which one ERGM is used to represent the partnership formation process, and another (potentially different) model to represent the dissolution process. In the conditional logit expression, the formation model is specified as:

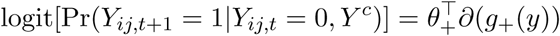

where now the edge *Y_ij_* is indexed by time and formation at time *t* + 1 is conditional on no edge *Y_ij_* existing at time *t*. Network statistics and coefficients are unique to the formation model. The analogous form for the dissolution model is:

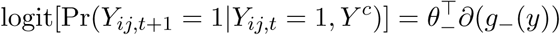

where the edge dissolution is in fact mathematically expressed and solved in terms of its compliment, edge persistence conditional on current edge existence.

#### Data for estimating TERGMs

Dynamic network models may be estimated from several different forms of empirical data: panel data from a network census over time, a single snapshot of a cross-sectional network census with additional data on partnership duration, and an *egocentrically sampled* crosssectional network with duration data. For egocentric network sampling, a random sample of the population is drawn, and respondents are asked about a subset of their partnerships within a relevant time interval (Morris 2004). The partners are not recruited into the sample. The statistical theory for estimating ERGMs from egocentrically sampled data can be found in Krivitsky and Morris (2017). The dynamic data structures and algorithms are described in more detail in the documentation for the **tergm** and **networkDynamic** packages (Krivitsky and Handcock 2015; Butts *et al*. 2016).

**EpiModel** was designed to work with cross-sectional egocentrically sampled network data with retrospective information on relational persistence. While this is the most limited form of network data, it is also the form most likely to be available in many epidemiological contexts. In an egocentric study, one can measure network summary statistics such as the distribution of the number of ongoing partnerships (the momentary degree distribution), broken down by nodal attributes, selective mixing patterns in nodal attributes at the dyad level, the age of active partnerships, and the duration of completed partnerships. The network models in **EpiModel** are based on TERGMs that are estimated from these network summary statistics. Note that these summary statistics may also be observed from a network census, along with a much wider range of additional statistics (e.g., triads and larger cycles), and **EpiModel** can also use these if they are available. Examples of how egocentric network data are translated into network statistics then used for epidemic modeling are provided in detail in our applied modeling papers (Goodreau *et al*. 2010, 2012; Jenness *et al*. 2016b,a).

### 3.3. Core modeling functions

**EpiModel** simulates epidemics over dynamic networks by integrating the TERGM framework for the estimating and simulating the partnership process with other modules that stochastically simulate the processes related to infection and demographics. Network epidemic models require specification of the parameters for each of these processes, and for the relationships among them.

#### Process dependence

The three processes represented in **EpiModel** – network, epidemic, and demography – are unique, but possibly dependent, steps in the modeling process. The key issue is whether the partnership network dynamics are *independent* of or *dependent* on the other processes over time. Network structure always influences the epidemic process, so the question is whether the epidemic influences the network structure. Examples of such feedback are when behavior depends on disease status, or when infection induces mortality. All open population models imply dynamics (e.g., birth, death or migration) that change the network with respect to the set of nodes, so models with these demographic processes always induce network dependence.

*Independent process models* assume no influence of epidemics on the network structure, and no demographic dynamics. Under this assumption, one can simulate the entire time series of the dynamic network first, and then run the epidemic process on top of this dynamic (but structurally fixed) network. This is computationally very efficient, and potentially useful for pedagogical and other demonstration purposes, as it simplifies the interpretation of the simulated epidemic dynamics. But it is only realistic for a limited set of real-world epidemic models: asymptomatic infections (since any infection that is symptomatic may induce behavior change) or short time frames (where demography can safely be ignored).

*Dependent process models* instead allow the epidemic and demographic processes to influence the network structure. Computationally, this means that the network has to be simulated one step at a time, alternating with the simulation and updating of the epidemic and demographic processes. This approach is clearly required for infections like HIV, where infection status may influence partner selection (referred to in the HIV literature as “seroadaptive behavior,” since it is a blood-borne pathogen). It is also required for all open population models that allow for births, deaths, and migration. One of the benefits of using TERGMs is that they provide a simple closed-form adjustment to account for the impact of changing population size and composition on partner selection (Krivitsky *et al*. 2011). Thus, the network model does not need to be re-estimated, the coefficients are simply updated at each time step to reflect current epidemic and demographic conditions. **EpiModel** has been designed to handle these adjustments automatically.

## 4. Base models

Simulating an epidemic on a network in **EpiModel** requires three core functions:

1. `netest` estimates the generative model for the dynamic partnership networks. This function calls the `ergm` and `stergm` functions from the **statnet** `R` packages. `netest` includes several features for fitting temporal ERGMs specifically within the epidemic modeling framework, discussed in our examples below.
2. `netdx` simulates one or more replications of the dynamic network over time from the model fit in `netest`. This is used to run diagnostics on whether the fitted model reproduces the empirically observed network summary statistics. Diagnostics are a critical step in the modeling sequence to ensure that the final output can be interpreted properly.
3. `netsim` runs the stochastic epidemic processes over a network simulated from `netest`. For independent models, the entire dynamic network time series is simulated first, followed by epidemic simulation over this dynamic network realization. For dependent models, the simulation sequentially updates the network, the disease transmission, and the demography at each time step; this allows for full dependence among these processes, but is computationally slower than an independent model of comparable length.

Full specification of a base model in the network class involves setting the initial demographic conditions, specifying the formation and dissolution models and target statistics for the network, selecting the structure of the epidemic model (SI, SIS, SIR) and setting its initial conditions and parameters. Below, we provide two examples of base network models in **EpiModel**, one independent and one dependent.

### 4.1. Example 1: Independent SIS model

Before we get started, we set the random number seed to allow for reproduction of the exact results below, since there are stochastic algorithms behind the model estimation and simulation processes. In the accompanying `R` file, we redo this before starting each of the four examples.

~~~
*R> set.seed(12345*)
~~~

In this example, we model a curable disease where recovery does not induce immunity (e.g., a bacterial STI) and infection status does not influence behavior, in a closed population. Heterogeneity in the nodes will be represented by a two-level risk group designation, with some assortative mixing by level. In network terminology, this is a *one-mode* network, because ties are allowed between all node types.

The values we use in this example are chosen for convenience; in a research setting, these would instead be empirically derived from data. Section 6 discusses the ongoing challenges in model parameterization.

#### Step 1: Estimating network structure

The population is set up as a network from the beginning. We start with an empty network (i.e., one with no ties) containing 1000 persons in two equally sized risk groups. The `network.initialize` function creates the empty network and the `set.vertex.attribute` function adds the risk group attribute, which is a binary variable that will be used to control infection risk as a function of network structure.

~~~
*R> nw <- network::network.initialize(n = 1000, directed = FALSE*)
*R> nw <- network::set.vertex.attribute(nw, "risk", rep(0:1, each = 500)*)
~~~

The two `R` functions listed above are contained in the **network** package. These will be the only functions in this tutorial directly calling another **statnet** package, and therefore we have used the :: reference to make this explicit (even though it is not technically necessary because **EpiModel** loads **network**).

#### Network model estimation and diagnostics

The network formation and dissolution formulas specify how persons in the population form and dissolve partnerships over time. The `formation` object is a right-hand side `R` formula object that follows the terminology and methods of the **ergm** and **tergm** packages (Handcock *et al.* 2017; Krivitsky and Handcock 2015). All terms are defined in the `"ergm-terms"` help file, which can also be accessed in the help documentation for those packages.

Here we will include terms to set the following network properties for the formation model:

1. *Density:* this sets the baseline probability of forming a tie, and is typically included in all models (analogous to an intercept term in a linear model). The ERGM term for this is `edges`. The statistic is the count of the number of edges (partnerships) in the network.
2. *Heterogeneity by attribute:* this allows the probability of an edge to depend on the nodal attribute “risk” we set up above. In our model, we will use it to set a higher mean degree for a high-risk group. The ERGM term for this is `nodefactor`. The statistic is the count of the number of ties for members of each attribute value. Note that each tie will count twice, once for each node.
3. *Selective mixing by attribute:* this allows the probability of an edge to depend on the attributes of both nodes. It is used to capture the level of assortative (or disassortative) mixing between groups. The ERGM term for this is `nodematch`. The statistic is the count of the number of ties between nodes of the same risk group. Here a tie will count only once.
4. *Degree distribution:* there are many ways to specify further heterogeneity in the degree distribution. In the absence of further specification, the conditional probability of a partnership forming is Bernoulli with the (group-specific) parameter determined by the coefficients on the terms above, and the resulting degree distribution is a Binomial mixture. Here we will modify this by specifying the number of nodes who have more than one partnership at any time. The ERGM term for this is `concurrent`. The statistic is the count of the number of nodes with two or more ties at any time step.

The resulting formation model is specified as the following ERGM formula:

~~~
*R> formation <- ~edges + nodefactor("risk") + nodematch("risk") + concurrent*
~~~

With egocentric inference the data are passed to the ERGM estimation function in summary form as *target statistics* for each term. If we had data, we could calculate these target statistics from them. Here, for this example, we will pass target statistics with fabricated values that are internally consistent:

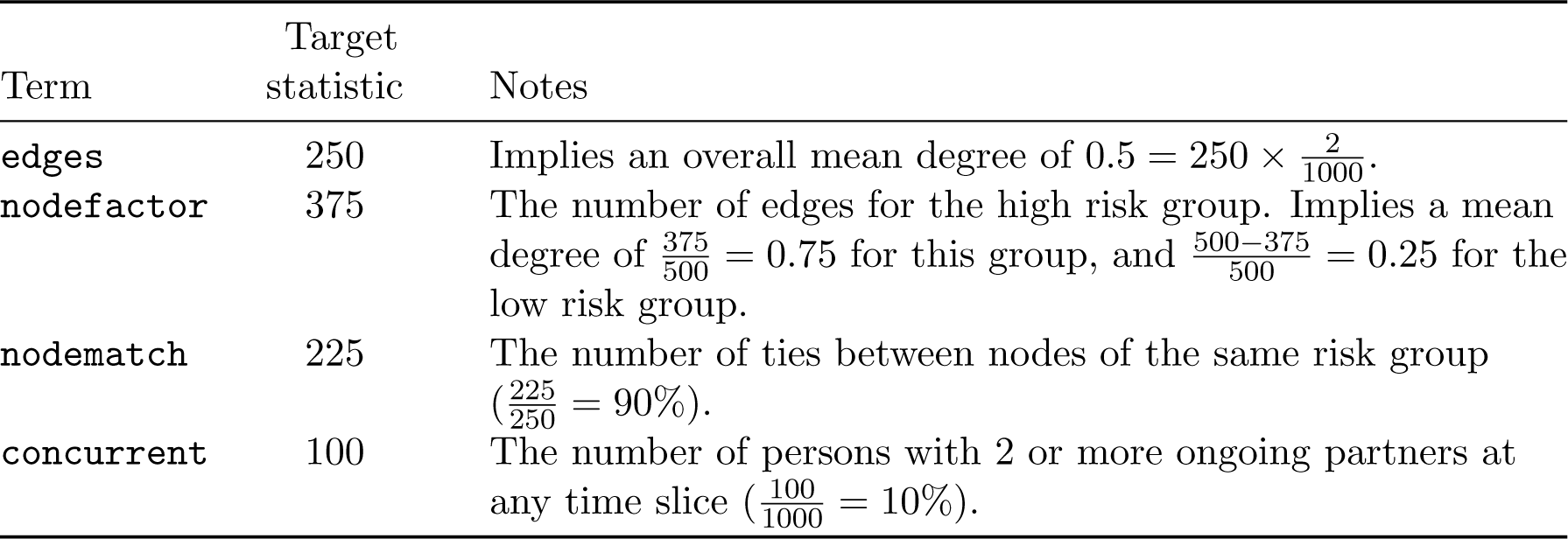

| Term | Target statistic | Notes |
| --- | --- | --- |
| **edges** | 250 | Implies an overall mean degree of $0.5 = 250 \times \frac{2}{1000}$. |
| **nodefactor** | 375 | The number of edges for the high risk group. Implies a mean degree of $\frac{375}{500} = 0.75$ for this group, and $\frac{500-375}{500} = 0.25$ for the low risk group. |
| **nodematch** | 225 | The number of ties between nodes of the same risk group ($\frac{225}{250} = 90\%$). |
| **concurrent** | 100 | The number of persons with 2 or more ongoing partners at any time slice ($\frac{100}{1000} = 10\%$). |

Note that for the nodefactor term, the default reference category is the lowest value of the factor (here 0, the low-risk group), so target stats are only provided for the comparison factor levels (here, only level 1, the high-risk group); the targets for the default group are set implicitly by the edges term. In addition, for this statistic, each tie counts twice, once for each node, so the denominator is twice the number of ties.

~~~
*R> target.stats <- c(250, 375, 225, 100*)
~~~

These target statistics represent the expected values of the network statistics for each crosssectional slice of the dynamic network time series.

A critical factor to consider when using target statistics to specify an ERGM formula is that these statistics are not independent, and they must be logically consistent. For example, the count of partnerships within the same risk group (the `nodematch` target statistic), cannot logically be higher than the total number of partnerships in the population (the edge target statistic). As another example, setting the edge target statistic to some lower value, say 50 instead of 250, has implications for other degree-related target statistics. An edges count of 50 would imply a mean degree of 0.1. Under a Poisson distribution, the expected proportion of persons with a degree greater than 1, given that mean degree, would be less than 1% (in `R, ppois(1, 0.1, lower.tail = FALSE)` calculates this). Fitting an ERGM with a much lower mean degree but keeping the same target for concurrency of 10% is nearly mathematically impossible; when this happens the MCMC-based estimation methods will fail unless the concurrency target is also adjusted downward.

**EpiModel** will generate error messages if the model fitting fails, but the content of these messages is produced by the deeper underlying packages, and not always entirely clear. Informative error messages from poorly fit models is an ongoing area of software development for both **EpiModel** and **statnet**. Here, it is challenging due to the many possible combinations of formulas and target statistics. This is one reason that we emphasize that these models are most useful when driven with empirical data, from a probability sample. In that case there is a strong chance of logical consistency. However, consideration in model specification, including experimentations with the dependencies within ERGM terms and target statistics associated with them, is a key part of the learning process for network modeling. Our general advice is to start simple, building complexity incrementally and only when necessary.

For the dissolution model in **EpiModel**, we take advantage of a computational shortcut to calculate the dissolution model coefficients analytically as a function of the duration of the ties (Carnegie *et al*. 2014). These coefficients are then passed into the dissolution model as fixed *offset* terms (an offset is a model parameter that is fixed by design, not estimated).

In this example, we specify a dissolution model with an edges term, meaning that the probability of edge dissolution at each time is a homogeneous constant hazard across partnerships, and therefore does not depend on the specific configuration of persons within that partnership. The resulting distribution of partnership durations, therefore, is geometrically distributed as we are working in discrete time. Other dissolution models (heterogeneous by nodal attributes) are supported by **EpiModel** (see the help file for `dissolution_coefs`), while other hazard specifications (e.g., duration dependent) are an area of ongoing development.

If we had empirical data, we would estimate the mean age of active ties and use that here. For this example, we specify an example mean duration of 80 time steps; this implies, at any time step, each partnership has a 1.25% risk of dissolving.

~~~
*R> coef.diss <- dissolution_coefs(dissolution = ~offset(edges), duration = 80*)
*R> print(coef.diss*)
Dissolution Coefficients
=======================
Dissolution Model: ~offset(edges) Target Statistics: 80
Crude Coefficient: 4.369448
Mortality/Exit Rate: 0
Adjusted Coefficient: 4.369448
~~~

The output from this function indicates both adjusted and crude coefficients, which are equivalent in this case. When working with open populations, where persons leave the network through death or migration they will differ, as we will see in the next example.

The `netest` function calls the estimation routines from **tergm** (Krivitsky and Handcock 2015) to estimate the coefficients for the formation and dissolution model. Inputs are the base network, the formation formula, the formation target statistics, and the dissolution coefficient.

~~~
*R> est1 <- netest(nw, formation, target.stats, coef.diss*)
~~~

By default, **tergm** exploits a computational efficiency when possible by using an approximation that relies on the assumption that prevalence = incidence *×* duration. When this assumption holds, dividing the coefficients estimated from a cross-sectional ERGM fit to the target statistics (which represent configuration prevalence) by their corresponding dissolution offsets will approximate the dynamic formation coefficients (Carnegie *et al*. 2014) (the configuration incidence). The approximation is most accurate when time steps are short relative to partnership lengths; as a rule of thumb the approximation bias is minimal when the edge duration is greater than 20 time units and the mean degree is less than 1. The bias is also greater in highly structured dyadic-dependent models. This default approach can only be used if the dissolution model is a subset of the formation model.

#### Step 2: Diagnosing network fits

After the model fit completes, it is crucial to diagnose the output, especially when one is relying on the approximation method. There are two forms of model diagnostics for a dynamic ERGM fit with `netest`: static and dynamic. When the approximation method has been used, static diagnostics check the fit of the cross-sectional ERGM to the target statistics. Dynamic diagnostics check the fit of the model adjusted to account for edge dissolution. Both function by simulating networks from the fitted model and comparing summary statistics from the simulated networks to the observed targets. When running a dynamic network simulation as we do with **EpiModel**, it is good to start with the dynamic diagnostics, and if there are fit problems, work back to the static diagnostics to determine if the problem is due to the cross-sectional fit itself or with the dynamic adjustment (i.e., the approximation method).

Here we will examine dynamic diagnostics only. These are run with the `netdx` function, which simulates from the model fit object returned by `netest`. We simulate from the model 10 times over 1000 time steps. The values are chosen based on the stochasticity in the model, which is a function of network size, model complexity, and other factors. In general, choosing these control parameters (number and length of simulations) may be an iterative process; the only downside to choosing larger values is computational burden. With respect to the length of the simulations (number of time steps), this should be set to a much higher value than the mean edge duration specified for the model. By default, the network statistics that are diagnosed are those in the formation formula, but the `nwstats.formula` argument may be used to monitor any additional statistics of interest.

~~~
*R> dx <- netdx(est1, nsims = 10, nsteps = 1000*)
~~~

Printing an object output from netdx will show summary tables of the simulated network statistics against the target statistics. The `mean` and `sd` values for the formation diagnostics are the mean and standard deviations of those statistics across all simulations and time steps. In our example results below, the simulated edges mean is slightly higher than the target, but within a standard deviation. The difference may be a result of either random variation (for which more simulations may be needed) or systematic bias primarily stemming from the use of the estimation approximation described above (for which further diagnostics described below would be necessary).

~~~
*R> print(dx*)
EpiModel Network Diagnostics
=======================
Diagnostic Method: Dynamic
Simulations: 10
Time Steps per Sim: 1000
Formation Diagnostics
~~~

|  | Target | Sim Mean | Pct Diff | Sim SD |
| --- | --- | --- | --- | --- |
| edges | 250 | 251.788 | 0.007 | 14.505 |
| nodefactor.risk.1 | 375 | 379.114 | 0.011 | 24.771 |
| nodematch.risk | 225 | 226.374 | 0.006 | 13.481 |
| concurrent | 100 | 101.861 | 0.019 | 10.899 |

|  | Target | Sim Mean | Pct Diff | Sim SD |
| --- | --- | --- | --- | --- |
| Edge Duration | 80.000 | 72.708 | -0.091 | 71.411 |
| Pct Edges Diss | 0.012 | 0.013 | 0.005 | 0.007 |

There are two forms of dissolution diagnostics. The edge duration row shows the mean age of partnerships across the simulations; it tends to lie below the target since the mean includes a burn-in period where all edges start at a duration of zero (illustrated below in the plot). To use this as a diagnostic for matching edge duration, a very long simulation interval is needed. The next row shows the percent of current edges dissolving at each time step, and is not subject to bias related to burn-in. The percentage of edges dissolving is the inverse of the expected duration: if the duration is 80 time steps, then we expect that 1/80 or 1.25% dissolve each day. The standard deviation for dissolution is close to the mean, as is expected, since our model of memory-less relational dissolution implies a geometric distribution for relational length, and this is a feature of that distribution.

Plotting the diagnostics object will show the time series of the target statistics against their targets (the black dashed lines). The solid lines show the mean values of the statistics across all simulations, and the bands show the inter-quartile range of values (the range of the bands may be changed with the `qnts` argument in `plot.netdx`). Similar to the numeric summaries, the plots in Figure 2 show a good fit over the time series.

**Figure 2:**
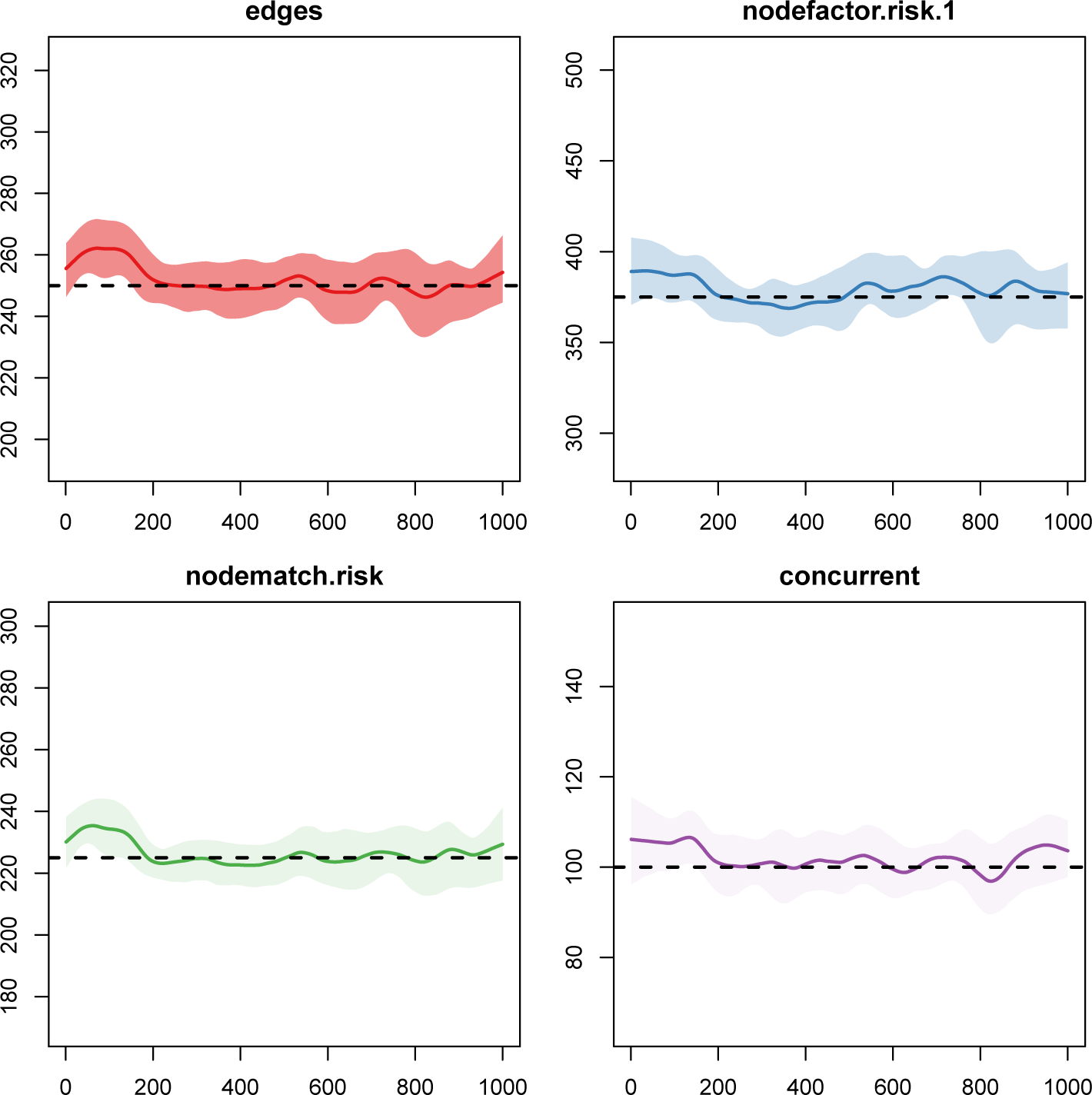
Formation model diagnostics.

~~~
*R> plot(dx*)
~~~

The dissolution model fit is plotted with either a `duration` or `dissolution` type, as defined above. The effect of the burn-in on the duration statistic is clear in the first plot, as the duration hits the target value of 50 after about 200 time steps. Both metrics indicate a good fit of the dissolution model to the target, as shown in Figure 3.

**Figure 3:**
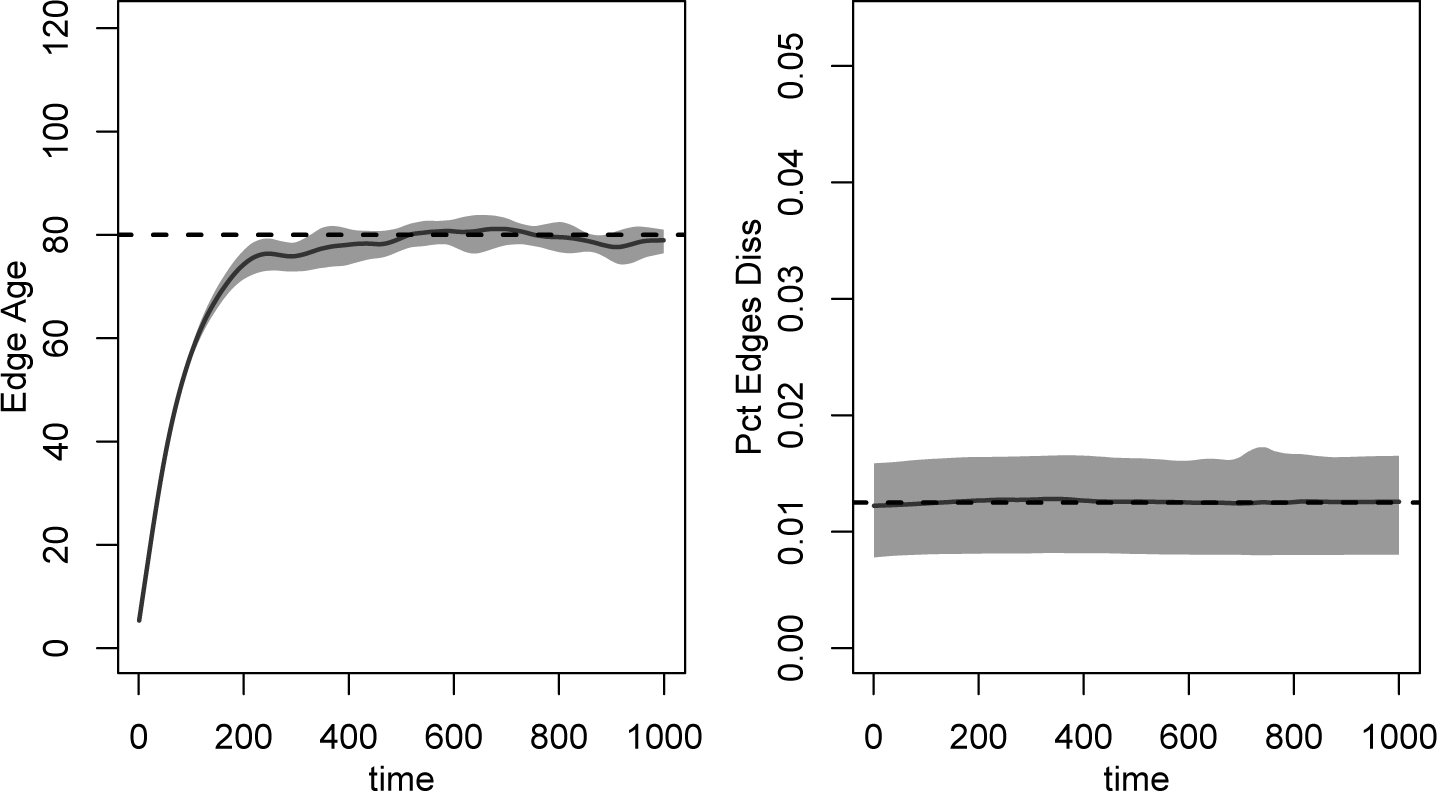
Dissolution model diagnostics.

~~~
*R> par(mfrow = c(1, 2)*)
*R> plot(dx, type = "duration"*)
*R> plot(dx, type = "dissolution"*)
~~~

If the model diagnostics had suggested poor fit, then additional diagnostics and fitting would be necessary. If using the approximation method, one should first start by running the crosssectional diagnostics (see the `dynamic` argument in `netdx`). If the cross-sectional model fits well but the dynamic model does not, then a full STERGM estimation (not using the approximation method mentioned above) may be necessary; this can be done with the argument `edapprox=FALSE`. If the cross-sectional model does not fit well, different control parameters for the ERGM estimation may be necessary (see the help file for `netdx` for instructions).

#### Step 3: Epidemic model setup and simulation

**EpiModel** simulates epidemics over dynamic networks in the independent process case by simulating the epidemiological processes such as infection transmission and disease recovery on the dynamic network that was simulated above. These processes also have stochastic elements so that the range of potential outcomes under the model specifications can be observed. **EpiModel** uses three helper functions to input epidemic parameters, initial conditions, and other control settings for the epidemic model.

For initial conditions, we use the `i.num` argument to set the initial number infected at the start (a more precise allocation of infection onto specific nodes can be made using the `status.vector` argument). For simplicity, this example will use a starting prevalence of 5%, equally distributed over risk groups.

~~~
*R> init <- init.net(i.num = 50*)
~~~

The base SIS model requires three parameters. The *infection probability* (`inf.prob)` is the risk of transmission per act between a susceptible person and an infected person. The *act rate* (`act.rate`) is the mean number of acts that occur within each active partnership during each time step. The mean frequency of acts per person per time step is the mean number of partnerships per person multiplied by this act rate parameter. The *recovery rate* (`rec.rate`) is the probability that an infected person recovers at a given time step. In an SIS model, they become susceptible again after recovery.

These three disease-related parameters are set using the helper function `param.net`:

~~~
*R> param <- param.net(inf.prob = 0.1, act.rate = 5, rec.rate = 0.02*)
~~~

The value used for the recovery rate implies that the mean duration of infection is 50 time steps.

The control settings contain the structural features of the model, which include the epidemic type, number of time steps per simulation, and number of simulations. The `epi.by` argument allows us to pass in a categorical nodal attribute which will be used to stratify the prevalence outcome statistics.

~~~
*R> control <- control.net(type = "SIS", nsteps = 500*,
*+ nsims = 10, epi.by = "risk"*)
~~~

Once all of the necessary model specifications have been set, simulating the model is straight-forward. We pass the fitted network model object along with the epidemic parameters, initial conditions, and control settings to the netsim function.

~~~
*R> sim1 <- netsim(est1, param, init, control*)
~~~

#### Examining model output

Printing the model output lists the inputs and outputs of the model. The output includes the sizes of the compartments (`s.num` is the number susceptible and `i.num` is the number infected) and flows (`si.flow` is the number of infections and `is.flow` is the number of recoveries). The output may be extracted using the `as.data.frame` function, which we do not show here but is described in its documentation page.

~~~
*R> print(sim1*)
EpiModel Simulation
=======================
Model class: netsim
Simulation Summary
-----------------------
Model type: SIS
No. simulations: 10 No. time steps: 500 No. NW modes: 1
Model Parameters
-----------------------
inf.prob = 0.1
act.rate = 5
rec.rate = 0.02
Model Output
-----------------------
Variables: s.num s.num.risk0 s.num.risk1 i.num i.num.risk0
i.num.risk1 num num.risk0 num.risk1 is.flow si.flow
Networks: sim1 … sim10
Transmissions: sim1 … sim10
~~~

Time-specific epidemic statistics may be obtained using the summary function. This summary shows the mean and standard deviation of simulations at time step 500.

~~~
*R> summary(sim1, at = 500*)
EpiModel Summary
=======================
Model class: netsim
Simulation Details
-----------------------
Model type: SIS
No. simulations: 10
No. time steps: 500
No. NW modes: 1
~~~

**Figure 4:**
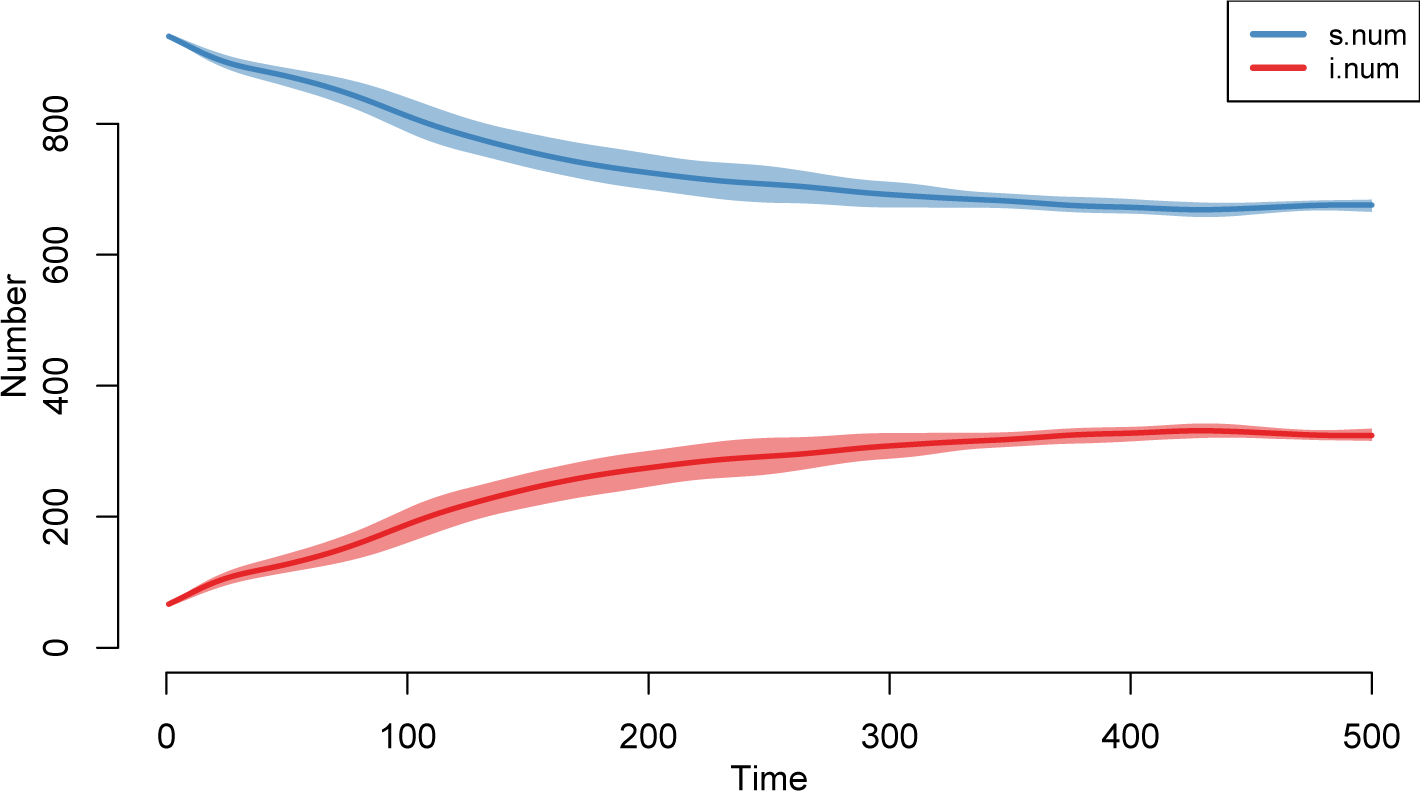
Epidemic model default plot.

|  | mean | sd | pct |
| --- | --- | --- | --- |
| Suscept. | 675.9 | 17.233 | 0.676 |
| Infect. | 324.1 | 17.233 | 0.324 |
| Total | 1000.0 | 0.000 | 1.000 |
| S -> I | 5.8 | 2.486 | NA |
| I -> S | 5.5 | 3.064 | NA |

The default plot for a netsim object, shown in Figure 4, will display the prevalence of the compartments in the model across simulations. The means across simulations are plotted with the thicker lines and the polygon band shows the inter-quartile range across simulations. Individual simulation lines are also available; consult the help page for `plot.netsim` for plotting parameter options.

~~~
*R> plot(sim1*)
~~~

To plot the incidence of infection and recovery, we specify these outcome variables using the y argument. As shown in Figure 5, the two lines converge at approximately time step 300, which is consistent with the stabilization of prevalence at that time as shown in the prevalence plot.

**Figure 5:**
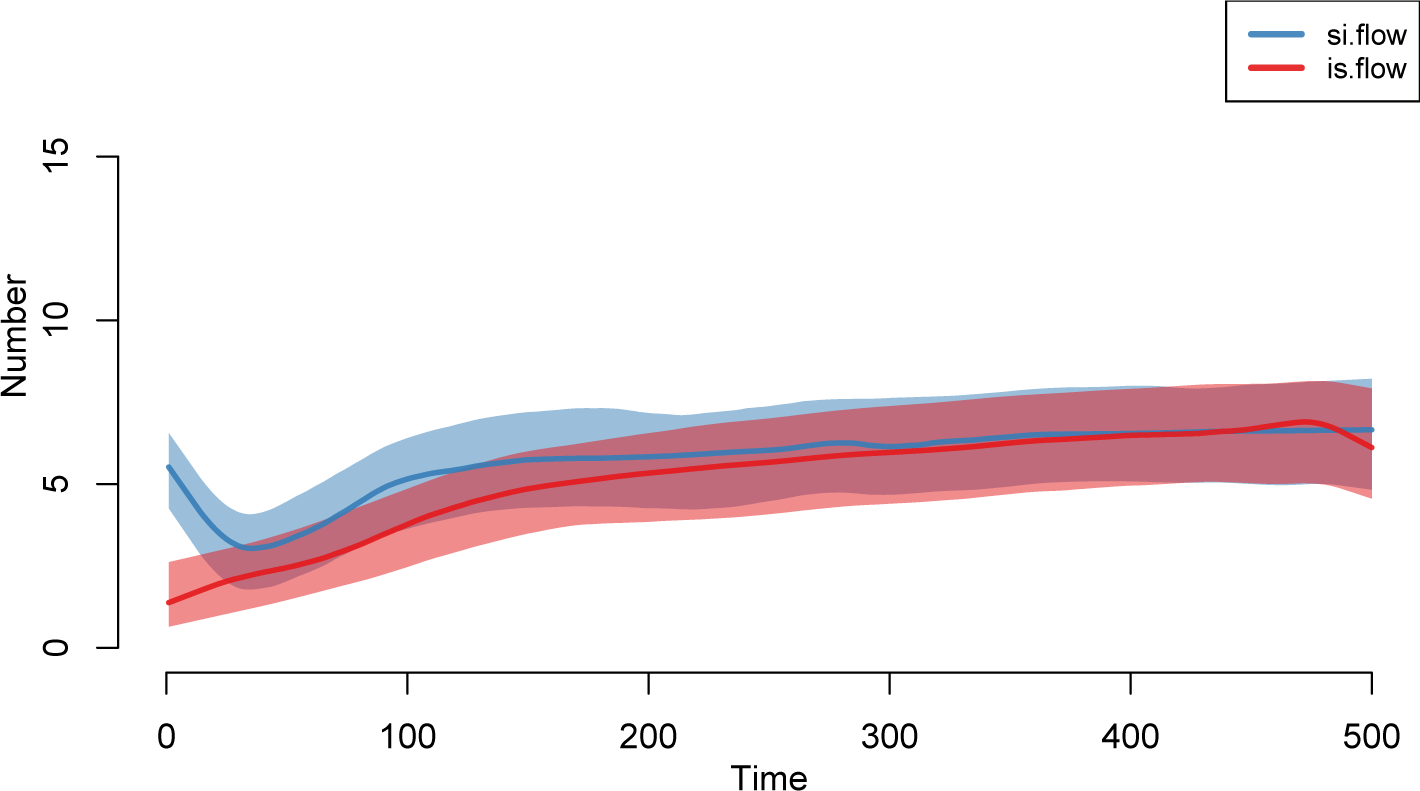
Incidence and recoveries.

~~~
*R> plot(sim1, y = c("si.flow", "is.flow"), leg = TRUE*)
~~~

To compare the results by risk group, the stratified outputs are passed to y. Figure 6 shows that the prevalence of disease is much higher for the high-risk group as a function of their higher mean degree and the containment of the epidemic within that group as a result of the high assortivity.

**Figure 6:**
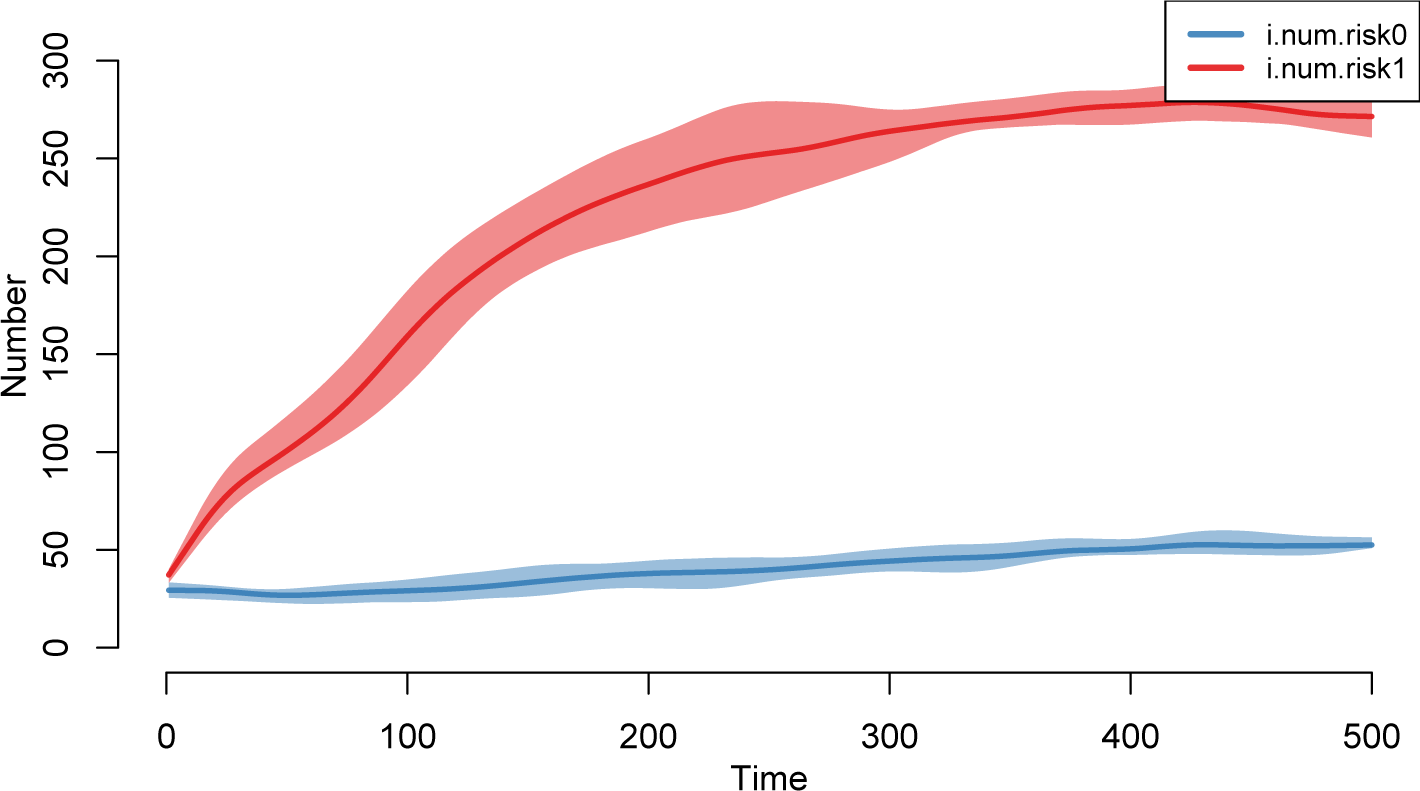
Stratified results.

~~~
*R> plot(sim1, y = c("i.num.risk0", "i.num.risk1"), leg = TRUE*)
~~~

Plotting the static network at different time steps and over different simulations can show the patterns of partnership formation and infection spread over those partnerships. The network plot is available by specifying type="network". In Figure 7 we plot two time points from among the simulations that are closest to the mean across all simulations with respect to disease prevalence, at time steps 1 and 500. We chose the mean here to easily illustrate a *representative* simulation; other options are the specific simulation number (1 to 10), "max", and "min" (see the help documentation for `plot.netsim`). The `col.status` argument handles the color coding for easy plotting of the infected (red) versus susceptible (blue).

**Figure 7:**
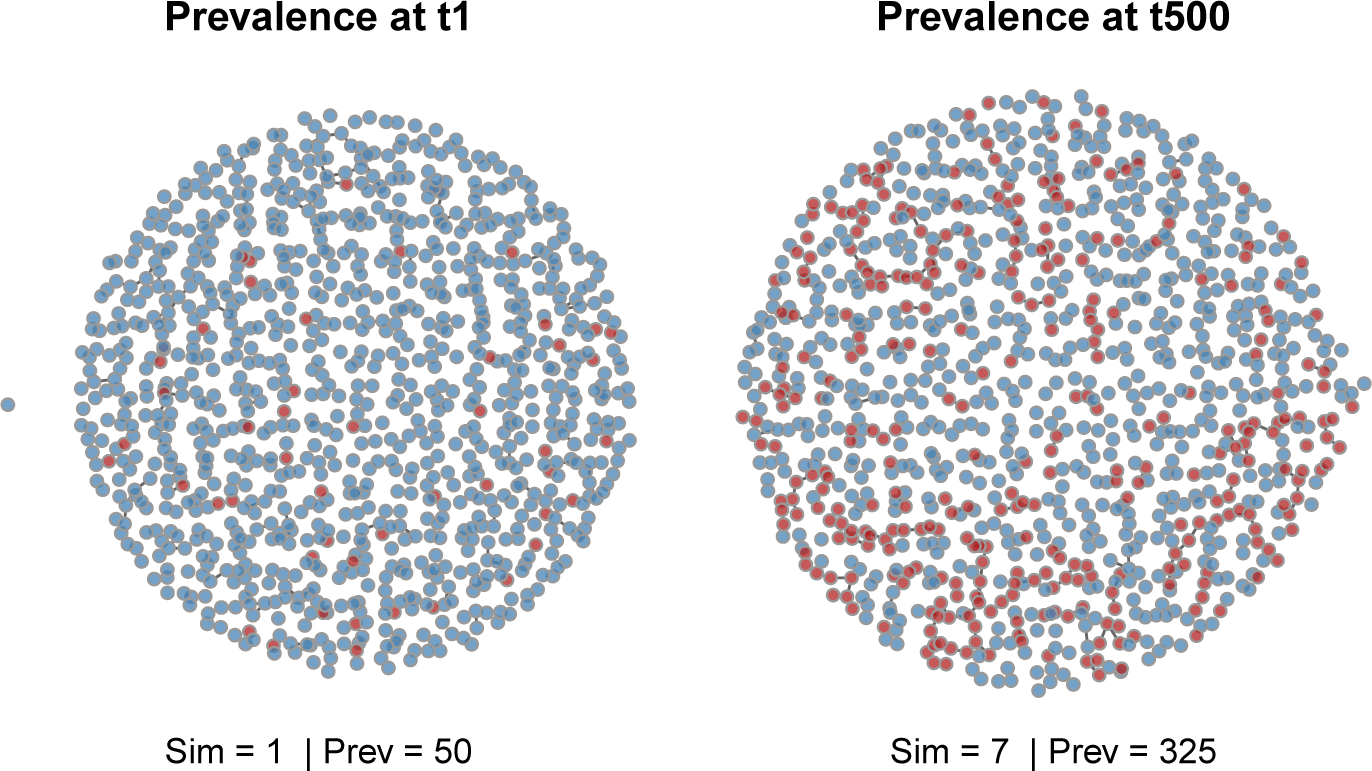
Static network plots.

~~~
*R> par(mfrow = c(1,2), mar = c(0,0,1,0)*)
*R> plot(sim1, type = "network", at = 1, sims = "mean"*,
*+ col.status = TRUE, main = "Prevalence at t1"*)
*R> plot(sim1, type = "network", at = 500, sims = "mean"*,
*+ col.status = TRUE, main = "Prevalence at t500"*)
~~~

### 4.2. Example 2: Dependent SI model

In this example, we allow the network to be influenced by demographic dynamics, and the demographics to be influenced by the epidemic process.

There are three types of process dependence built into the base models within the stochastic network class in **EpiModel**:

1. **Demographic vital dynamics:** Births and deaths (or more generally, entries and exits) will always influence the network because the exit of persons removes their associated ties, and entry creates new persons (initially without partners) with whom ties may be formed. Vital dynamics also influence the epidemic process by replenishing susceptibles and removing infecteds.
2. **Infection-adaptive behavior:** Infection status can influence the probability of partnership formation and dissolution, as well as behavior within partnerships. Thus, it can influence both the network model and the epidemic process.
3. **Infection-dependent vital dynamics:** Infection status can also influence fertility and mortality, allowing the epidemic to feed back on the demographic processes.

In each of these cases, changes over time in the node set and/or infection status require the network to be updated before simulating each step.

#### Adjusting the network model for vital dynamics

In the presence of vital dynamics, two adjustments must be made to the coefficients of the temporal ERGM: adjusting the formation model for the impact of changes in network size and composition, and adjusting the dissolution model for the impact of death/exit on tie removal. The former occurs automatically in the **tergm** software; the latter is executed using a helper function in **EpiModel**.

When the network size (number of nodes) changes, the ratio of ties to nodes changes. As a result, either network density or the mean degree will change; both cannot be preserved at the same time, since mean degree is approximately equal to *N × density*. This raises the question of what the appropriate time-invariant parameterization should be for a network model. Preserving density, for example, implies that someone who moves from a village of 500 to a city of 50,000 would increase the number of ties they have by a factor of 100. In practice, most social or sexual relationships of interest to epidemic modelers do not behave this way; they instead preserve mean degree when network size changes.

The default (canonical) parameterization of ERGMs generally preserves density for dyadic-independent models in the context of changing network size, with some more complex behavior for dyadic-dependent models (see (Kolaczyk and Krivitsky 2015)). But ERGMs are also easily adjusted to preserve mean degree instead Krivitsky *et al*. (2011), and **EpiModel** is designed with this behavior as the default. Adjustment involves modifying the edges coefficient, *θ*, in the formation ERGM:

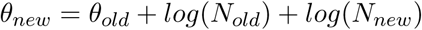

where *N_old_* is the old population size and *N_new_* is the new population size. This adjustment is handled within netsim in **EpiModel** automatically, although if the user prefers to preserve density in response to population change (or some intermediate between density and mean degree), that may be toggled with arguments in the `control.net` helper function.

Partnerships are subject to two forces of dissolution: endogenous (e.g., break-ups or divorces) and exogenous (the death of one or both partners). The STERGM model, when estimated from empirical data that implicitly includes both processes, captures the effects from both processes. If we then layer an additional mortality process on top of the network during the epidemic simulation, our simulated dissolution rate will be too high; relational durations become shorter than observed and network density becomes lower. The solution is to adjust the dissolution parameter in the STERGM to approximate the endogenous dissolution process only. The dissolution_coefs function automatically applies a *death correction* adjustment to account for the competing risks, storing the relevant information in a way that the subsequent components of the simulation can use. In the simple offset(edges) homogeneous dissolution model, we derive that through conditional probabilities. Starting with the observation that the probability of an edge dissolving at time *t*, *P* (*E_t_*), is:

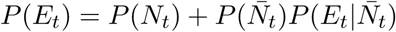

where *P* (*N_t_*) is the probability that either node dies (which automatically dissolves an edge), 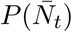 is probability that both nodes survive, and 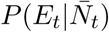 is the probability that the edge dissolves endogenously (i.e., given that both nodes survive). This formula may be rearranged to solve for that endogenous dissolution probability:

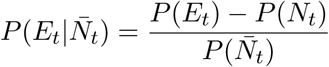

Since the dissolution coefficient, *θ*, is expressed in terms of edge persistence (1 *− P* (*E_t_*)), and on the log-odds scale, we calculate the corrected *θ* as:

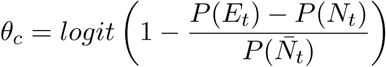

Note that there are many different approaches for handling the competing risks like these within discrete-time simulations, but this approach works correctly under standard assumptions in our epidemic simulations. This adjustment is demonstrated directly in the example below.

#### Network initialization

In this example, we will simulate an epidemic model on a bipartite network. Bipartite network structures are used when all ties must be *between* groups, called **modes**. This structure may be used to model, for example, STI transmission in a strictly heterosexual population. **EpiModel** provides some built-in utilities that facilitate mode-specific network terms (e.g., sex-specific degree distributions) and epidemic parameters (e.g., transmission asymmetries).

As before, the first step is to specify a network structure, including features like size and nodal attributes. Here, we construct an empty network of 1000 nodes, with 500 in each mode. Although not explicitly named as such, we conceive of the first mode as females and the second mode as males.

~~~
*R> num.m1 <- num.m2 <- 500*
*R> nw <- network::network.initialize(num.m1 + num.m2, bipartite = num.m1, + directed = FALSE*)
~~~

In this example, we will also set up a degree distribution for each sex that might be obtained from an egocentric sample. We need to ensure that the number of partnerships implied by the women’s degree distribution is the same as that implied by the men’s. This does not require that the distributions are the same, just the total number of partnerships.

Our example population has an even 1:1 sex ratio. We will set the mean proportion of females and males having 0, 1, 2, or 3 partners at any one time as follows:

~~~
*R> deg.dist.m1 <- c(0.40, 0.55, 0.04, 0.01*)
*R> deg.dist.m2 <- c(0.48, 0.41, 0.08, 0.03*)
~~~

Note that men report more concurrency (having greater than one partner) than women do, but more men than women also report having no partners.

In **EpiModel**, we use the check_bip_degdist function to check that the implied number of edges matches across modes, given the degree distributions and the mode sizes set above:

~~~
*R> check_bip_degdist(num.m1, num.m2, deg.dist.m1, deg.dist.m2*)
~~~

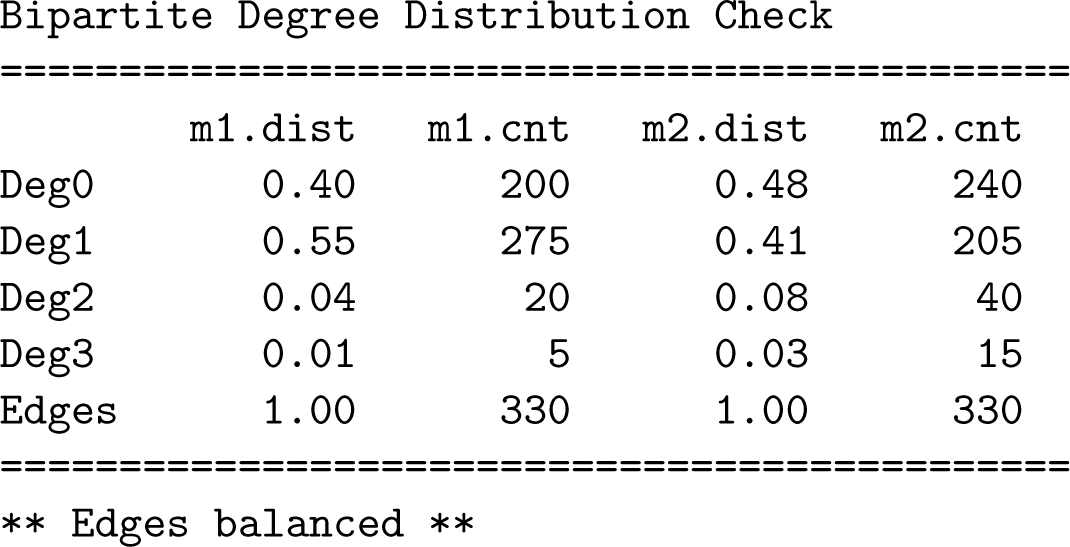

The table shows the fractional distribution and number of nodes with each degree, as well as the total number of edges implied by each degree distribution. Here the total edges for females and males match, and we are good to proceed. In other cases, one may encounter unbalanced numbers when using egocentric data for a variety of reasons (e.g., sampling, reporting errors), even though any bounded empirical population with two modes would in theory have to balance if a census were taken. When faced with this, network modelers must consider how they wish to address this (e.g., by averaging the two modes’ mean degree somehow) before proceeding (see Morris (1991); Koehly *et al.* (2004) for examples).

#### Network model estimation and diagnostics

In this example, we will use the formation model to specify details of the sex-specific degree distributions, not just the mean degree.

The ERGM terms b1degree(d) and b2degree(d) can be used to specify a non-parametric model for the degree distributions for a bipartite network, for modes 1 and 2 respectively. Here, we will fit the network statistics for the sex-specific degrees of 0 and 1, leaving the upper tails of the degree distributions for each sex free to vary. The choice of which terms to include in any ERGM formula such as this depends on the research question of interest, as well as the statistical degrees of freedom available in potentially highly correlated terms. In this case, the mode-specific network size and the total number of edges remove available degrees of freedom, such that we effectively have a fully specified model. In general, it is best to start with a less specified model and build up terms to add further complexity as necessary.

~~~
*R> formation <- ~edges + b1degree(0:1) + b2degree(0:1*)
~~~

As in the first example, we will input the data in the form of target statistics from our initial network for each of these terms. These can be extracted directly from the bipartite check table above.

~~~
*R> target.stats <- c(330, 200, 275, 240, 205*)
~~~

The adjustments to the formation model for the demographic dynamics is automatically handled by **tergm** during estimation (Krivitsky and Handcock 2015).

The dissolution model is a homogeneous constant hazard with mean partnership duration of 25 time units. Since there will be births and deaths in this model, we also need to specify an overall death rate, using the `d.rate` parameter of dissolution_coefs, to adjust for the competing risk of death on edge dissolution. Since the coefficient for the dissolution model represents the probability of edge persistence, the resulting value of the adjusted coefficient will be higher than the crude coefficient.

~~~
*R> coef.diss <- dissolution_coefs(dissolution = ~offset(edges)*,
*+ duration = 25, d.rate = 0.006*)
*R> print(coef.diss*)
Dissolution Coefficients
=======================
Dissolution Model: ~offset(edges)
Target Statistics: 25
Crude Coefficient: 3.178054
Mortality/Exit Rate: 0.006
Adjusted Coefficient: 3.533444
~~~

The value 0.006 was calculated as a weighted mean of the mortality rates specified as epidemic parameters below.

The netest function estimates the model as before.

~~~
*R> est2 <- netest(nw, formation, target.stats, coef.diss*)
~~~

Model diagnostics should be run before moving on to epidemic simulation, but one important caveat is that the networks simulated in netdx here do not include the demographic processes that could influence the structure of the network within the full epidemic model. So it will be necessary to also run *post-simulation* diagnostics after the epidemic model is simulated with netsim.

~~~
*R> dx <- netdx(est2, nsims = 10, nsteps = 1000*)
~~~

The `plots.joined` argument for `plot.netdx` places all the plots in one panel instead of a separate panel for each statistic. The simulations are all on target as shown in Figure 8; as before, the dashed lines show the target statistics, the solid lines are the values of the means of the simulated statistics over the 10 simulations, and the bands are the inter-quartile range of those same simulations.

**Figure 8:**
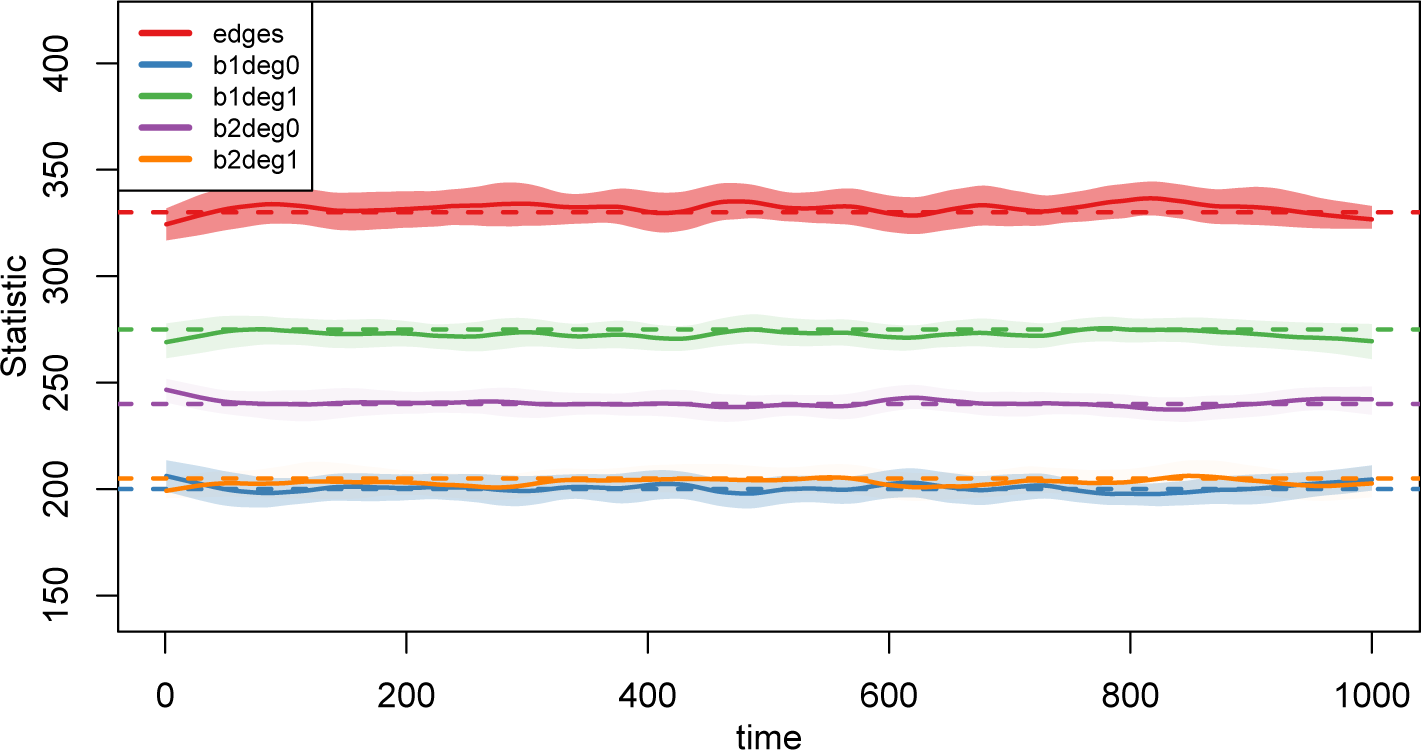
Network diagnostics plot.

~~~
*R> plot(dx, plots.joined = TRUE*)
~~~

#### Epidemic model setup and simulation

We set the initial number infected to 50 for both the first and second modes.

~~~
*R> init <- init.net(i.num = 50, i.num.m2 = 50*)
~~~

The parameters for both infection and demographic processes are set here, since the demographic rates may be influenced by infection state. In this example, we will use different transmission probabilities by mode, with a three-fold higher susceptibility of infection for females compared to males. The birth rates are parameterized with the `b.rate.m2` parameter set as NA to signify that the number of births into the second mode (males) should be based on the size of the female population. The mortality rate is specified so that females have an overall lower death rate than men, and that there is an excess risk of mortality upon infection.

~~~
*R> param <- param.net(inf.prob = 0.3, inf.prob.m2 = 0.1*,
*+ b.rate = 0.006, b.rate.m2 = NA*,
*+ ds.rate = 0.005, ds.rate.m2 = 0.006*,
*+ di.rate = 0.008, di.rate.m2 = 0.009*)
~~~

In the control settings, the `delete.nodes` argument removes inactive (i.e., deceased) nodes from the network object at each time step for computational efficiency. The `nwstats.formula` argument, similar to its definition in the `netdx` function, allows monitoring of a set of network statistics that may differ from the formation formula. Here, we explicitly monitor the mean degree with the `ergm` term `meandeg`.

~~~
*R> control <- control.net(type = "SI", nsims = 10, nsteps = 500*,
*+ nwstats.formula = ~edges + meandeg*,
*+ delete.nodes = TRUE*)
~~~

Once the model specifications are complete, the simulation is again straightforward.

~~~
*R> sim2 <- netsim(est2, param, init, control*)
~~~

#### Examining model output

With dependent processes, the coefficients of the network model are updated at the end of each time step to reflect changes in the size of the network; the network at the next time step is then simulated with the updated coefficients. When using base models, this adaptive simulation happens automatically whenever certain parameters are included in the function calls, such as rates for vital dynamics.

Because of these changes, it is important to examine the network structure from the simulations before moving on to interpreting the epidemic outputs. One particularly important check is to see that the vital dynamics processes did not lead to any systematic biases in mean degree, given that this quantity is our time-invariant target and is influenced by several approximations and adjustments, and given how fundamental it is to transmission dynamics. The fact that we had the algorithm calculate this as the simulation unfolded in the previous command makes this straightforward. Here, we also add a target line for expected mean degree manually, since it is not an explicit term in the model formulas.

~~~
*R> plot(sim2, type = "formation", plots.joined = FALSE*)
*R> abline(h = 0.66, lty = 2, lwd = 2*)
~~~

Figure 9 shows that the number of edges declined substantially over time, but the mean degree was preserved at its target of 0.66. The decline in edges was a result of the overall declining population size, which is apparent from the next plot. However, the adjustment to maintain mean degree has worked as intended.

**Figure 9:**
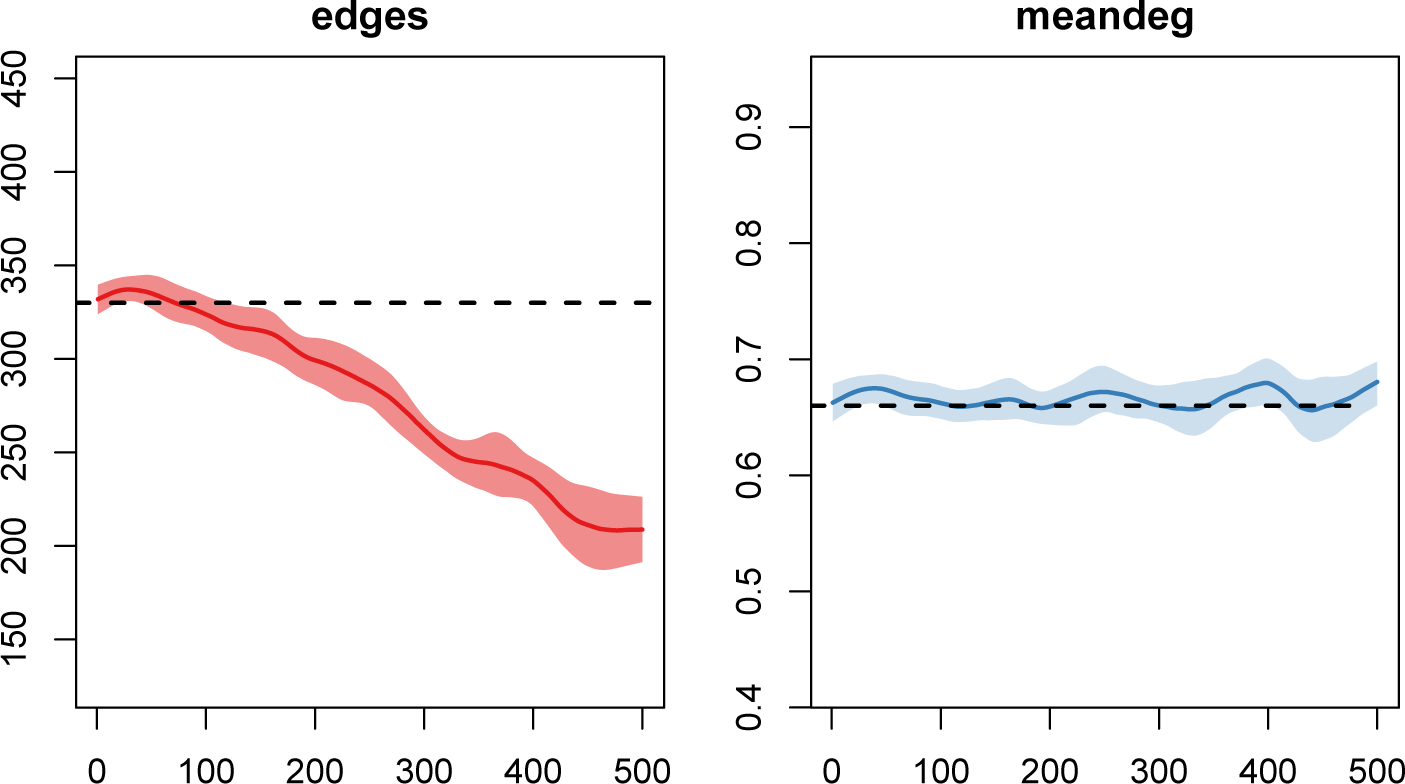
Post-simulation diagnostics.

The default plot of the simulation object shows the means of the compartment sizes over time for each of the modes (quantile bands are suppressed by default for bipartite simulations).

~~~
*R> par(mfrow = c(1,2), mar = c(3,3,1,1), mgp = c(2,1,0)*)
*R> plot(sim2, popfrac = TRUE*)
*R> plot(sim2, popfrac = FALSE*)
~~~

The left panel of Figure 10 shows the prevalence of infection states (by mode, if the model includes them), whereas the right panel shows the absolute numbers in each compartment. The overall population size is declining substantially due to disease-induced mortality, which has an effect on both sexes in both disease states.

**Figure 10:**
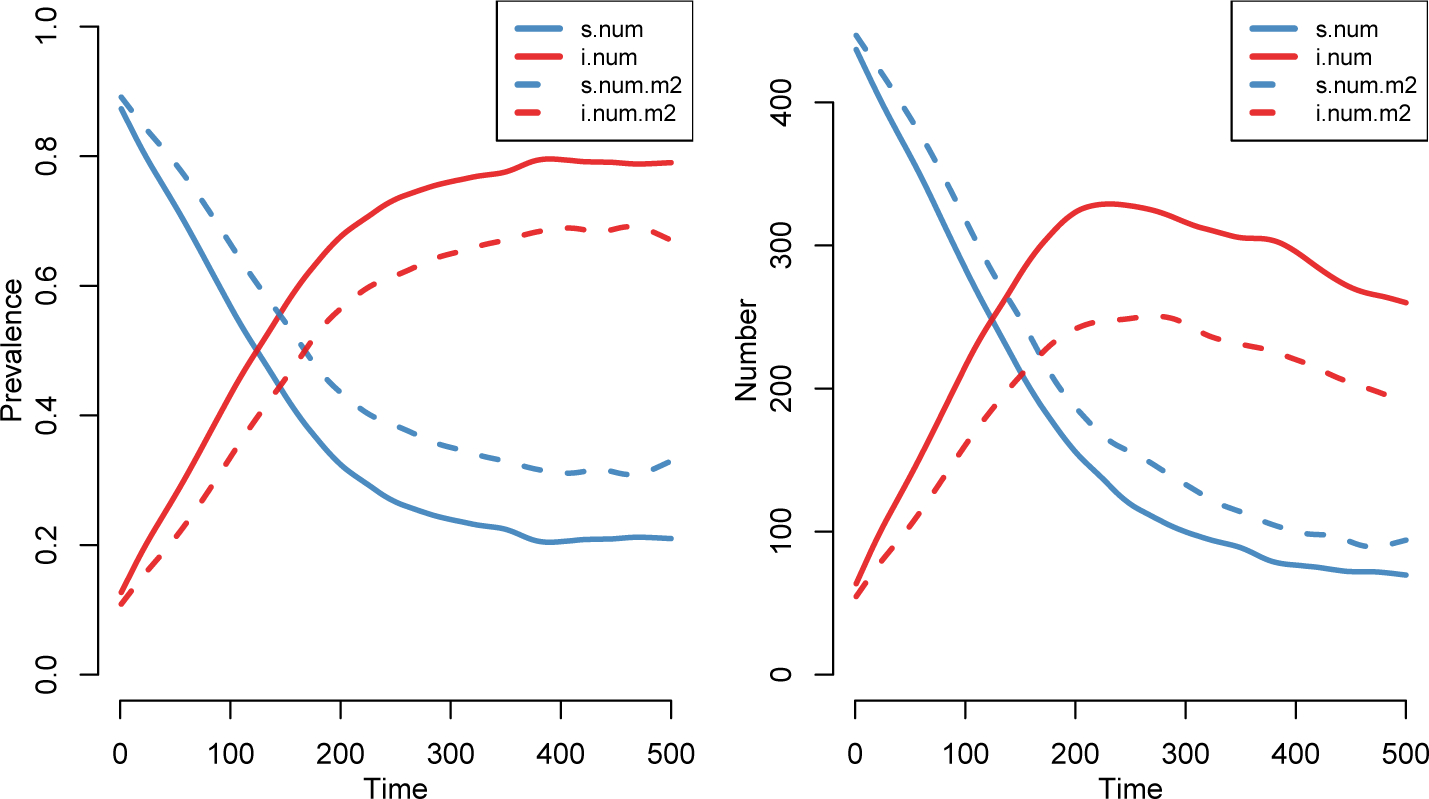
Prevalence and absolute epidemic size.

## 5. Model extension API

The models described so far – what we have called *base models* – are aimed at beginning **EpiModel** users. They are useful primarily for teaching and learning the basic principles of modeling epidemics on networks. For most applied modeling research, these base models will not have the range and flexibility to address specific pathogen, population, and intervention related questions. However, we have designed **EpiModel** so that it can be extended to incorporate new functionality for research purposes.

In this section, we describe **EpiModel**’s flexible application programming interface (API) for writing and simulating new network-based epidemic models for research. We provide two examples to demonstrate how to replace existing modules (e.g., substituting a new transmission module) or supplement the existing modules with new modules that control novel system mechanics.

Extending **EpiModel** involves writing new **modules** that are plugged into the simulation workflow executed by `netsim`. These modules will typically be run at each time step to simulate one or more of the processes that occurs within the larger system. Modules may depend on one another: a mortality module may reference an age attribute defined and updated by an aging module.

Research-level epidemic models tend to be quite complex, with many parameters and interactions between all of the processes. Understanding and validating the code involved in simulating systems is a central challenge in mathematical modeling. That also holds true for developing **EpiModel** extensions. But one need not start from scratch: a research model may combine existing model code as a starting template.

### 5.1. Modular extension framework

When running base models of the network class in **EpiModel**, the settings specified in `control.net` control a series of modules that govern how persons in the population become infected, recover, die, and so on. The recovery module, for example, is controlled by the `recovery.FUN` parameter in control.net. For each built-in module, there is a default function that defines the module. The default recovery function (as described in the help documentation for `control.net`) is `recovery.net`. When the time loop runs the series of specified time steps, this function is called and the individuals within the population are subject to the recovery processes defined within it.

Arguments in `control.net` that have a `.FUN` suffix are pointers that tell the main simulation function, `netsim`, to evaluate them. Each call to `netsim` begins by initializing the simulation under the conditions specified in `initialize.FUN` (with the default function `initialize.net`). Any of these default functions may be replaced by writing and specifying a new function that handles the same process. New modules that handle other types of processes not in the base model workflow may be added into the mix by including those module names ending in `.FUN`. New modules may be plugged into `control.net` because it contains a flexible … argument that handles new inputs of arbitrary definition.

To develop new modules, either to replace existing ones or to add to the set, one must understand the basic *rules* for module structure. The `netsim` function allows for broad flexibility in module definition, but components of the general structure must be followed. There are four core elements to the **EpiModel** API for epidemic network model modules:

1. The main data structure storing the information passed among the modules is `dat`, which is an `R` object of class list. All the inputs, such as parameters, and the outputs, such as summary epidemiological statistics, are contained in `dat`. Each module reads `dat`, updates internal data within `dat`, and then outputs `dat` for the next module to work on. As a result, every module has only two input arguments: `dat` and at, the current time step.
2. Attributes of individual nodes are stored in a sublist to `dat` called `attr`. This is a named list: each element corresponds to one attribute stored as a vector, and all vectors have the same length (the number of individuals in the population). netsim creates five attributes by default: `active, status, infTime, entrTime`, and `exitTime`. The `active` attribute keeps track of whether an individual is currently alive or has exited the population through some process such as death or out-migration. The `status` attribute indicates current infection status as a character (`s, i, r`, for susceptible, infected, and recovered in the base models); `infTime` records the time at which each individual was infected (`NA` for susceptible nodes); in the case of SIS models this reflects the most recent time of infection. The `entrTime` and `exitTime` attributes contain the time steps at which the person enters or leaves the network. Each attribute is model-specific and may be used in different ways to simulate different diseases (e.g., the `status` attribute would never be set to `r` if simulating SIS epidemics in which there is no permanent recovered state).
3. Summary model output statistics are contained in a sublist to `dat` called `epi`. This list stores information on the current *prevalence*, the number of nodes in each infection state, and *incidence,* the sizes of the flows between infection states. The default for base models is to calculate summary prevalence statistics in the `get_prev.FUN` module in the last module for each time step; summary incidence statistics are typically calculated in the module where the event occurred.
4. In **EpiModel** time loops, the initial network is coded as time 1. The only module that is run at this time step is the `initialize.FUN` module to set up components like network structure and infection status. All other functions will start running at time 2 and continue for the number of time steps specified in `control.net`. The choice of time step 1 to represent the initial network and 2 as the first step of the dynamic process simulation, while not traditional, is adopted in order to make data storage and querying align with `R`’s convention of indexing vectors beginning with 1, not 0. That is, disease prevalence at time 400, say, is now stored in position 400 of the prevalence vector, not in position 401. In the examples below, some new vectors are created at time step 2 and updated after that by using a shortcut: `if (at == 2) {create something} else {edit it}`. This creates a new data structure without needing to explicitly edit the initialization module. The latter can be unwieldy since it handles many tasks.

The examples below provide practice with this functionality. You can also explore these features and more by reading through the code for `netsim` and for the other functions that it calls, or by stepping through a call to `netsim` using a browser tool in `R` like `debug`.

### 5.2. Example 1: Age-dependent mortality extension

In this example, we show how to construct an aging process, age-specific mortality, and a constant growth model for births. This will require one *new* module function and two *replacement* module functions, along with the associated parameters.

#### New aging module

To introduce aging into a model, we need to write a new module function that we will call `aging` to perform the necessary processes. Writing this illustrates some key requirements of any module added to `netsim`.

At the first time step in the loop, which is step 2 for the reason noted above, persons in the population are randomly assigned an age between 18 and 69 years (uniformly distributed) in units that are congruent with our time steps. We might choose this age range because we collected data only on adults within this age group. Here, our simulation time step will be a month, so we specify age to the nearest month. At each subsequent time step, age is incremented by one month.

~~~
*R> aging <- function(dat, at) {*
*+ if (at == 2) {*
*+ n <- sum(dat$attr$active == 1*)
*+ dat$attr$age <- sample(seq(from = 18, to = 69 + 11/12, by = 1/12)*,
*+ n, replace = TRUE*)
*+ } else {*
*+ dat$attr$age <- dat$attr$age + 1/12*
*+ }*
*+ if (at == 2) {*
*+ dat$epi$meanAge <- c(NA, mean(dat$attr$age, na.rm = TRUE)*)
*+ } else {*
*+ dat$epi$meanAge[at] <- mean(dat$attr$age, na.rm = TRUE*)
*+ }*
*+ return(dat*)
*+ }*
~~~

Note that the new module, as required, has the two functional arguments `dat` and `at`. The `active` attribute stored in `dat` indicates whether each person is currently active, and it is evaluated at time step 2 to establish the size of the active population, `n`, so the `sample` function knows how many ages to produce. The new attribute `age` is stored as a vector in the `attr` list and updated for everyone at each time step. Any summary statistics we want to monitor can be stored in `dat$epi`. Here, we create a vector called `meanAge` at time step 2 for the current mean age of the active nodes in the population; the mean is updated and appended to the vector at each time step.

#### Modified death module

For this extension, we will replace the existing death module for susceptibles with a new one. In the existing module, the probability of death is based on a fixed risk that may vary by disease status; the parameters that control this, `ds.rate` and `di.rate` and so on, are set through `param.net`. Here, we will replace the module used by the base models with a new one that exploits our new age attribute. The probability of death will now be specified as a nonlinear function of increasing age:

~~~
*R> ages <- 18:69*
*R> death.rates <- 1/(70*12 - ages*12*)
~~~

Since we are using monthly time steps, the death rates are specified with that same scale. The rates could be estimated from life table data, but here we instead use a simple parametric form for age-specific mortality in which death rates are specified to increase monotonically with age (up to age 70). In our new death module, maximum age will be treated as a variable parameter, used to calculate the death rate in the module function. For each active person, death is then evaluated at each time step based on a draw from a Bernoulli distribution with probability equal to their age-specific risk.

~~~
*R> dfunc <- function(dat, at) {*
*+ idsElig <- which(dat$attr$active == 1*)
*+ nElig <- length(idsElig*)
*+ nDeaths <- 0*
*+ if (nElig > 0) {*
*+ ages <- dat$attr$age[idsElig]*
*+ max.age <- dat$param$max.age*
*+ death.rates <- pmin(1, 1/(max.age*12 - ages*12)*)
*+ vecDeaths <- which(rbinom(nElig, 1, death.rates) == 1*)
*+ idsDeaths <- idsElig[vecDeaths]*
*+ nDeaths <- length(idsDeaths*)
*+ if (nDeaths > 0) {*
*+ dat$attr$active[idsDeaths] <- 0*
*+ dat$attr$exitTime[idsDeaths] <- at*
*+ dat$nw <- deactivate.vertices(dat$nw, onset = at, terminus = Inf*,
*+ v = idsDeaths, deactivate.edges = TRUE*)
*+ }*
*+ }*
*+ if (at == 2) {*
*+ dat$epi$d.flow <- c(0, nDeaths*)
*+ } else {*
*+ dat$epi$d.flow[at] <- nDeaths*
*+ }*
*+ return(dat*)
*+ }*
~~~

The vector of IDs eligible for the process is determined by the current value of the active variable. The ages of those eligible are extracted from the `attr` list, the maximum age parameter is queried from the parameter list, `param` from `param.net`, and the death rate is calculated for each person based on their age and that maximum. A Bernoulli draw is performed for each eligible person with those rates, and if death results, then the active attribute for that person is toggled from 1 to 0.

The network object, located at `dat$nw`, is then updated with the changes. Each newly dead node is `deactivated` from the network. This process also removes their existing edges (if any) and prevents them from having future partnerships. A summary statistic vector is created for the number of deaths during each time step and saved as `d.flow` in the epi sublist.

#### Modified birth module

In this example, we will modify the birth module used by the base models. The new module replaces the base growth rate specification, a fixed percentage increase based on the active population at each time step, with a constant growth model. In this model, the size of the population is specified to grow *linearly* at each time step based on the rate parameter `growth.rate` (in contrast to exponential growth that typically occurs). The expected size of the population at a time step, *t*, is a function of the initial population size and this rate. Because nodes can also exit at each time step, the expected number of new births at each time step is the difference between the expected population size and the current population size. New births are added stochastically based on a Poisson distribution draw with the rate parameter equal to this difference.

~~~
*R> bfunc <- function(dat, at) {*
*+ growth.rate <- dat$param$growth.rate*
*+ exptPopSize <- dat$epi$num[1]*(1 + growth.rate*at*)
*+ n <- network.size(dat$nw*)
*+*
*+ numNeeded <- exptPopSize - sum(dat$attr$active == 1*)
*+ if (numNeeded > 0) {*
*+ nBirths <- rpois(1, numNeeded*)
*+ } else {*
*+ nBirths <- 0*
*+ }*
*+ if (nBirths > 0) {*
*+ dat$nw <- add.vertices(dat$nw, nv = nBirths*)
*+ newNodes <- (n + 1):(n + nBirths*)
*+ dat$nw <- activate.vertices(dat$nw, onset = at*,
*+ terminus = Inf, v = newNodes*)
*+ }*
*+ if (nBirths > 0) {*
*+ dat$attr$active <- c(dat$attr$active, rep(1, nBirths)*)
*+ dat$attr$status <- c(dat$attr$status, rep("s", nBirths)*)
*+ dat$attr$infTime <- c(dat$attr$infTime, rep(NA, nBirths)*)
*+ dat$attr$age <- c(dat$attr$age, rep(18, nBirths)*)
*+ }*
*+ if (at == 2) {*
*+ dat$epi$b.flow <- c(0, nBirths*)
*+ } else {*
*+ dat$epi$b.flow[at] <- nBirths*
*+ }*
*+ return(dat*)
*+}*
~~~

The value for the linear growth rate parameter is again passed to the module function through `param.net` and stored in `dat$param`. The expected population size uses the starting population size stored in the summary statistics list `epi`, and the current population size is calculated again with the `active` attribute in `dat`.

Any birth function must do two things to update the network: add new nodes to the network object and set all of their attributes with their initial values. In our function, the `add.vertices` function adds new nodes onto the network, and the `activate.vertices` function allows them to form edges. We also need to set the five default attributes and our new age attribute for the new nodes, on the `attr` list. Thus, new births will be active, susceptible to infection, have no infection time, have an entry time equal to `at`, have no exit time, and be age 18. The data for the new births is added to the end of the existing vectors.

#### Network model estimation

For the examples in this section, we will demonstrate how to specify the simplest possible population and network: a Bernoulli Random Graph (also known as an Erdös-Renyi graph). The nodes are homogeneous, and all ties have the same formation and dissolution probabilities, so partner selection is random and the degree distribution is binomial. The formation model only requires the target number of edges as input. For the dissolution coefficient adjustment, we take the mean of the unweighted age-specific death rates as a simple approximation of the overall mean death rate.

~~~
*R> nw <- network::network.initialize(500, directed = FALSE*)
*R> est3 <- netest(nw, formation = ~edges, target.stats = 150*,
*+ coef.diss = dissolution_coefs(~offset(edges), 60, mean(death.rates))*)
~~~

Since they were demonstrated above, we will skip the diagnostics here in the interests of brevity, but remind the reader that they should always be done after the network model has been estimated.

#### Epidemic model setup and simulation

To parameterize the epidemic model, it is necessary to collect all the new elements that were created into the `param.net` function. The transmission modules are unchanged in this example so they can use the same infection probability and act rate parameters as the base models. Our modified birth module requires a growth rate parameter. Since our time units are months, we specify a 1% yearly growth rate in terms of monthly growth. The modified death module requires an input of a maximum age, in years. The initial conditions can be specified like a base SI model.

~~~
*R> param <- param.net(inf.prob = 0.15, growth.rate = 0.01/12, max.age = 70*)
*R> init <- init.net(i.num = 50*)
~~~

For control settings, we specify the model type, number of simulations, and time steps per simulation, as we would for any base model. To replace the default death and birth modules, we specify that the existing `deaths.FUN` and `births.FUN` arguments take our new `dfunc` and `bfunc` functions respectively. Both of these functions must be sourced into memory before running `control.net`; the control function will search for them and save them within the object output.

To add the new aging module into `netsim`, we need to specify a new module name, followed by `.FUN` suffix so that **EpiModel** will register it as a module. The name can be anything, as long as it is not the name of an existing module. Here, we call our new module `aging.FUN`, and have it use the new `aging` function we wrote above.

~~~
*R> control <- control.net(type = "SI", nsims = 5, nsteps = 500*,
*+ deaths.FUN = dfunc, births.FUN = bfunc*,
*+ aging.FUN = aging, depend = TRUE*)
~~~

It is important to note the ordering of when the modules are executed within each time step. By default, all new modules like `aging.FUN` are run first and all existing (built-in) modules (modified or not) run second; new modules will be run in the order in which they appear in `control.net`, but built-in modules (modified or not) will be run in the order in which they are listed in the `control.net` help documentation. This ordering of all the modules may be changed and explicitly specified using the `module.order` argument in `control.net`.

Once the modules have been revised, all components are again simply passed to the netsim function.

~~~
*R> sim3 <- netsim(est3, param, init, control*)
~~~

#### Examining model output

Printing the model object shows that we now have non-standard birth and death output for flows. Additionally, we have the `meanAge` data saved as other output (summary statistics saved in `epi` named ending with `.num` will automatically be classified as compartments and `.flow` as flows).

~~~
*R> print(sim3*)
EpiModel Simulation
=======================
Model class: netsim
Simulation Summary
-----------------------
Model type: SI
No. simulations: 5
No. time steps: 500
No. NW modes: 1
Model Parameters
-----------------------
inf.prob = 0.15
growth.rate = 0.0008333333
max.age = 70
act.rate = 1
Model Output
-----------------------
Variables: s.num i.num num meanAge d.flow b.flow si.flow
Networks: sim1 … sim5
Transmissions: sim1 … sim5
~~~

Basic model plots in Figure 11 show the simulation results for the prevalence and absolute state sizes over time. All of the graphical plotting options available for base models can also be used for extended models.

**Figure 11:**
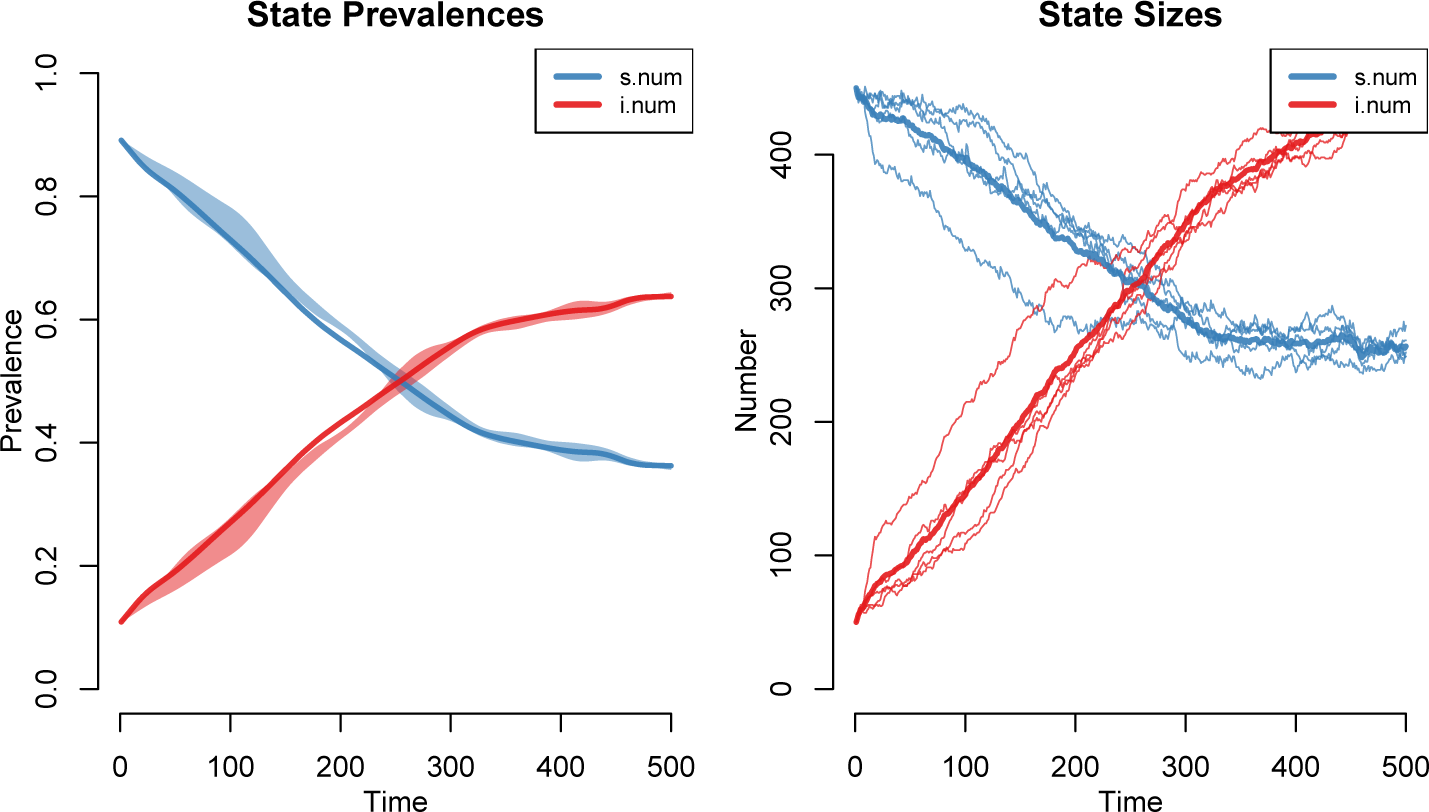
Extension epidemic model results.

~~~
*R> par(mfrow = c(1,2)*)
*R> plot(sim3, popfrac = TRUE, main = "State Prevalences"*)
*R> plot(sim3, popfrac = FALSE, main = "State Sizes", sim.lines = TRUE*,
*+ qnts = FALSE, mean.smooth = FALSE*)
~~~

This example demonstrated how to write modules using the **EpiModel** API to add system features not present in the base models. The API has been designed to make it relatively simple to modify existing modules or construct entirely new model elements, plug them into the software infrastructure, simulate from the new model, and analyze the results.

### 5.3. Example 2: SEIR epidemic model

In this final example, we will demonstrate how to modify the base disease types by adding a new state in the infection process: exposed but not yet infectious to others. This is traditionally referred to as an **SEIR** model, where the **E** compartment represents this initial stage of infection. Ebola is an example of a disease with this feature. In principle, this type of modification can also be used for adding any other relevant disease states to an SI/SIS/SIR model – like asymptomatic infection (e.g., for Chlamydia) or additional stages of infection (e.g., HIV and syphilis).

This example demonstrates a basic SEIR design with no births or deaths for the sake of simplicity. Similar to the previous example, we will have one replacement infection module and one entirely new module. Adding in an extra disease stage involves updating the infection module and adding a more generalized disease progression module that is based on the existing built-in recovery module.

#### Modified infection module

The built-in infection module is designed to handle a wide variety of specifications, most of which we will not use here, and to transition persons from a susceptible state directly to an infectious state. We will modify this, simplifying to remove unnecessary components and updating to represent the transition from susceptible to the new exposed but non-infectious state.

In this example, we will not print the full module code block, but it can be viewed in the `R` script file accompanying this article. The core structure of the module is exactly the same as the modules defined in the prior example, with operations occuring on the `dat` object before it is returned.

~~~
*R> infect <- function(dat, at) {*
*+*
*+ return(dat*)
*+ }*
~~~

The main processes in the built-in infection module are to extract a *discordant edgelist* – a matrix of ID numbers of active dyads in the network in which one member of the dyad is susceptible and the other is infected – and execute the stochastic process of transmission. Given epidemic parameters for the probability of infection per act (*τ*) and the number of acts per unit time (*α*), the per-partnership transmission rate is calculated using the standard equation 1 *−* (1 *− τ*)^*α*^. Transmission is a Bernoulli trial with this probability of infection.

~~~
*R> del <- discord_edgelist(dat, idsInf, idsSus, at*)
*R> del$transProb <- dat$param$inf.prob*
*R> del$actRate <- dat$param$act.rate*
*R> del$finalProb <- 1-((1-del$transProb)^del$actRate*)
*R> transmit <- rbinom(nrow(del), 1, del$finalProb*)
**R> del <- del[which(transmit == 1),]**
~~~

The key modification needed is to specify the newly infected person’s infection status as `"e"` rather than `"i"`.

~~~
**R> dat$attr$status[idsNewInf] <- "e"**
**R> dat$attr$infTime[idsNewInf] <- at**
~~~

The final lines within the module also need to be modified to track the size of this transition between susceptible and exposed states in a summary vector, `se.flow`, that will be available for analysis.

#### Construct a new disease progression module

The disease progression module will handle the transition from exposed to infectious, and also the transition from infectious to recovered in the model (so it will replace the built-in recovery module). The full structure of this module function, `progress`, is contained in the script file. Disease progression through each of these states is represented as an individual-level stochastic process, and unlike infection, there is a simple constant hazard of transition. The times spent in each disease state therefore follow a geometric distribution. The first part of the new module pulls two new parameters from the input lists that define the two rates of transition.

For each eligible person, the transition event is a Bernoulli trial with the rate parameter above. Persons who transition to the infectious state have their individual-level `status` attribute updated to the `"i"` value, at which point they are now capable of infecting others.

~~~
*R> nInf <- 0*
*R> idsEligInf <- which(active == 1 & status == "e"*)
*R> nEligInf <- length(idsEligInf*)
*R> if (nEligInf > 0) {*
*+ vecInf <- which(rbinom(nEligInf, 1, ei.rate) == 1*)
*+ if (length(vecInf) > 0) {*
*+ idsInf <- idsEligInf[vecInf]*
*+ nInf <- length(idsInf*)
*+ status[idsInf] <- "i"*
*+ }*
*+ }*
~~~

Persons who transition to recovered follow a process specified by the recovery function, which is simply copied from the existing recovery function into this new `progress` function.

The final job of this module is to calculate summary statistics and store them for analysis. The sizes of the susceptible and infected states are automatically calculated in all models. These states are contained in every epidemic model, so their summary statistics are calculated by default. In this module, we only need to handle the new exposed and recovered states:

~~~
*R> dat$epi$ei.flow[at] <- nInf*
*R> dat$epi$ir.flow[at] <- nRec*
*R> dat$epi$e.num[at] <- sum(active == 1 & status == "e"*)
*R> dat$epi$r.num[at] <- sum(active == 1 & status == "r"*)
~~~

#### Network model estimation

We again use a simple Bernoulli random graph model in order to focus on the new epidemiological inputs and structures.

~~~
*R> nw <- network::network.initialize(500, directed = FALSE*)
*R> est4 <- netest(nw, formation = ~edges, target.stats = 150*,
*+ coef.diss = dissolution_coefs(~offset(edges), 10)*)
~~~

We will again skip the diagnostics in the interests of brevity.

#### Epidemic model setup and simulation

There will be four variable parameters for the epidemic model, two of which are used in the infection module and the other two for the disease progression module. One must take care to ensure that the names of the variables are consistent between the inputs here and the module definition. In this example, we will set the mean time spent in the exposed state to half as long as in the infectious state.

~~~
*R> param <- param.net(inf.prob = 0.5, act.rate = 2*,
*+ ei.rate = 0.01, ir.rate = 0.005*)
*R> init <- init.net(i.num = 10*)
~~~

For the sake of simplicity, we will assume that the unit of time here is one day, so the average person in the population has 2 acts per day, spends 100 days in the pre-infectious exposed state (`1/ei.rate`), and 200 days in the infectious state before recovering (`1/ir.rate`).

For the control settings, the new infection function is passed to the existing `infection.FUN` argument, while the new progression module function is passed to a new `progress.FUN` argument. Since the disease progression function is handling the disease recovery process, the recovery module is toggled off by setting its function value to `NULL`. The `skip.check` setting toggles off some internal error checking intended for base models (e.g., that a `type` parameter is specified and the parameter values in `param.net` are consistent with the modules requirements).

~~~
*R> control <- control.net(nsteps = 500, nsims = 5, infection.FUN = infect*,
*+ progress.FUN = progress, recovery.FUN = NULL*,
*+ skip.check = TRUE, depend = FALSE, verbose.int = 0*)
~~~

Once the modules have been revised, all components are again simply passed to the *netsim* function.

~~~
*R> sim4 <- netsim(est4, param, init, control*)
~~~

#### Examining model output

The basic prevalence plot shows the outcomes of the four disease state sizes over time.

~~~
*R> par(mfrow = c(1, 1)*)
*R> plot(sim4, y = c("s.num", "i.num", "e.num", "r.num")*,
*+ mean.col = 1:4, qnts = 1, qnts.col = 1:4, leg = TRUE*)
~~~

We can see in Figure 12 that adding an exposed, non-infectious stage in the model has the effect of reducing the speed of the epidemic growth, but in this closed population nearly everyone gets infected within the simulation time frame for this parameter set.

**Figure 12:**
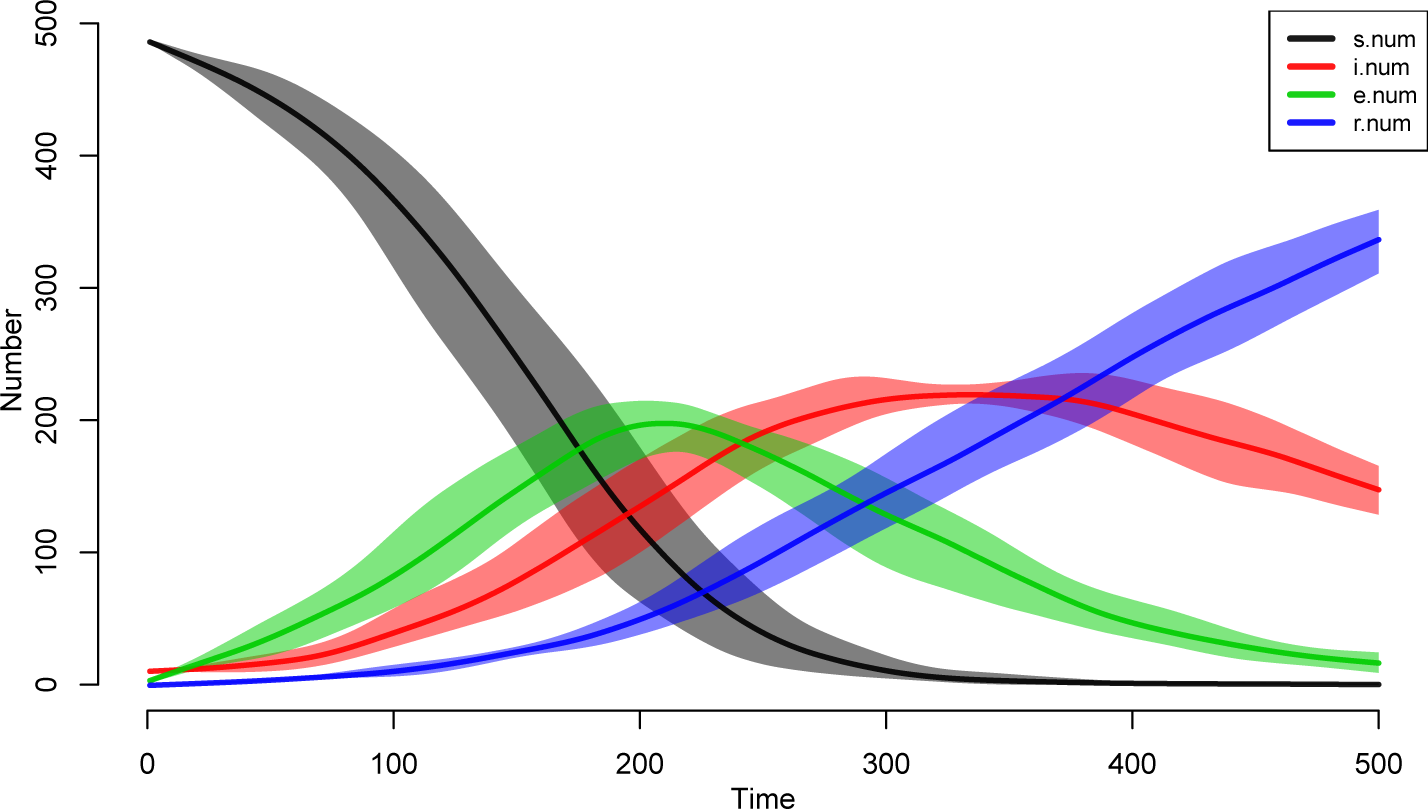
SEIR model output.

In summary, **EpiModel** provides a general, flexible, extendable framework for simulating epidemics over networks. The **EpiModel** API for extending the base models allows users to write new modules to control any aspect of the infection and demographic processes, and have these automatically feed back to the dynamic network structure. New modules are simply inserted into the model workflow that is executed by the `netsim` function. They are written as `R` functions (which may call functions in other languages such as C++), and the details of their execution are controlled directly as arguments to the `control.net` helper function.

## 6. Discussion

This paper has provided an overview of the **EpiModel** software platform for R. **EpiModel** is a tool for modeling infectious disease epidemics and other diffusion processes. **EpiModel** includes extensive tools and tutorials to simulate epidemics for three different classes of models (DCMs, ICMs, network models) with multiple modalities for each (Shiny apps, command-line programming). These tools may be used for teaching modeling, gaining insight into basic epidemic theory and phenomena, and developing novel research extensions.

The primary methodological contribution of **EpiModel** is the implementation of tools for modeling epidemics over dynamic networks based on Exponential-family Random Graph Models (ERGMs), a class of statistical models for network analysis that have been developed and supported within the **statnet** suite of packages for `R` (Handcock *et al*. 2015). In this paradigm, networks comprise a set of persons with (potentially overlapping) partnerships, where partnerships are defined as repeated disease-relevant contacts with the same person over time.

**EpiModel** provides seamless integration of the **statnet** functions for estimating and simulating complex network models, with a flexible set of functions for modeling the stochastic infection, recovery, demographic, and other processes that determine the trajectory of an epidemic. A key benefit of the **statnet** packages is that they have been designed to work with sampled network data and, in particular, egocentrically sampled network data. These data are relatively easy and inexpensive to collect, which makes these models practical for use in public health and other applied research contexts.

The EpiModel package provides a starting point for applied research. Most applications will need to make use of the flexible API to integrate the details of the pathogens, populations and intervention options that influence the specific transmission dynamics of interest. The package has been designed to support this kind of extension, informed, in part by our own research goals. This includes projects that focus on drivers of HIV infection in men who have sex with men (MSM) (Goodreau *et al*. 2012), network-related causes of racial disparities in HIV and STIs among heterosexuals in the United States (Morris *et al*. 2009), the role of acute-stage infection and concurrency for HIV in Zimbabwe (Goodreau *et al.* 2010), the combined impact of male circumcision and network structure on HIV in heterosexual couples in West Africa (Jenness *et al*. 2016b), the prevention benefits of combination HIV prevention among MSM globally (Sullivan *et al*. 2012), and the impact of new HIV prevention technologies among

MSM in the United States (Jenness *et al.* 2016a). Earlier projects relied on the **statnet** tools and prototyped code that led to the development of EpiModel, while the more recent projects have been programmed directly in **EpiModel** and led to the development of **EpiModelHIV**, an extension package of modules designed specifically to support modeling HIV transmission (available at https://github.com/statnet/EpiModelHIV). The open development platform on GitHub is intended to provide a mechanism for sharing such extensions to **EpiModel**.

In contrast to the basic model examples presented above, full-scale applied models require substantial literature synthesis and analysis to define model parameters. A central question in developing complex models like these is whether there are sufficient empirical data available for parameterization (Lessler and Cummings 2016). Each new layer of heterogeneity in the population or new transition between possible states in the model (e.g., disease status, clinical treatment, prevention interventions) should be grounded in realistic empirical data when available. Decisions to increase model complexity are often limited by available parameter data (Garnett *et al.* 2011). Bayesian statistical methods for dynamic models have been developed to estimate input parameters based on fitting these models to available epidemiological outcome data on disease prevalence and incidence (Poole and Raftery 2000). Approximate Bayesian Computation (ABC) is one approach we have used that is optimized for dynamic, stochastic models like ours (Toni *et al*. 2009). However, these robust methods are computationally complex, have identifiability problems as the number of parameters to be estimated is greater than the number of output statistics (as is typical), and require assumptions about the validity of the model structure conditional on the posterior parameter set.

Another related challenge in network modeling, present in microsimulation modeling generally, is the computational burden. Because stochastic models are simulated, typically over many iterations in large population sizes to obtain stable outcome measures, they require computational work greater than DCMs and other methods with analytic solutions. For

**EpiModel**, the computational burden is associated with both estimation and simulation. Estimation is usually relatively fast even for large network sizes because of the approximation method described above (Carnegie *et al.* 2014), but if this yields poorly fitting models then the full TERGM estimation may be required, which takes longer (Krivitsky and Handcock 2014). The resimulation of the network structure at each time step, accounting for changes to the node set (through vital dynamics) or composition of the nodal attributes, is the main computational bottleneck. In our applied network models, we often simulate networks with sizes of up to 25,000 nodes over 100 years or longer, depending on the research questions; these may take hours to compute depending on the number of simulations. We employ high-performance computing systems with many cores to conduct simulations in parallel to decrease the overall computing time. Ongoing work has been underway in both the **EpiModel** and **statnet** projects to further optimize the algorithms within the network resimulation in order to reduce this bottleneck. Those efforts will be described in the documentation for future releases of these packages.

We hope that the materials presented here provide a starting point for exploring the dynamics of epidemics on networks using **EpiModel**. There is a growing community of scholars interested in and using this software tool. Further resources, including links to an active mailing list, can be found the software website at http://epimodel.org/.

## 7. Acknowledgments

The authors would like to acknowledge the members of the **statnet** (Handcock *et al*. 2015) development team, including Mark Handcock, David Hunter, Carter Butts, Pavel Krivitsky, and Skye Bender-deMoll. Emily Beylerian, Skye Bender-deMoll, Kevin Weiss, and Li Wang have made important contributions to the **EpiModel** software code base. Kevin Weiss and

Emeli Anderson provided assistance in the review and editing of this paper. We thank the JSS editors and anonymous peer reviewers of earlier drafts of this work. The Network Modeling Group at the University of Washington has provided ongoing feedback on the software. Thanks also to students in our annual summer course, Network Modeling for Epidemics (course materials available at http://statnet.github.io/nme/), who have made suggestions and reported software bugs. This work has been funded by the United States National Institutes of Health Grants R01 HD068395, R21 MH112449, R21 HD075662, P30 DA027828, P30 AI050409, P30 AI027757, and T32 HD007543.

## References

Abu-Raddad L, Longini I (2008). “No HIV Stage is Dominant in Driving the HIV Epidemic in Sub-Saharan Africa.” AIDS, 22(9), 1055–61.

Anderson R, May R (1992). Infectious Diseases of Humans: Dynamics and Control. Oxford University Press.

Besag J (1974). “Spatial Interaction and the Statistical Anaylsis of Lattice Systems.” Journal of the Royal Statistical Society B, 36, 192–236.

Butts CT, Leslie-Cook A, Krivitsky PN, Bender-deMoll S (2016). networkDynamic: Dynamic Extensions for Network Objects. R package version 0.9.0, URL https://CRAN.R-project.org/package=networkDynamic.

Carnegie N, Krivitsky P, Hunter D, Goodreau S (2014). “An Approximation Method for Improving Dynamic Network Model Fitting.” Journal of Computational and Graphical Statistics, 24(2), 502–519.

Carnegie N, Morris M (2012). “Size Matters: Concurrency and the Epidemic Potential of HIV in Small Networks.” PLoS ONE, 7(8), e43048.

Chang W, Cheng J, Allaire J, Xie Y, McPherson J (2016). shiny: Web Application Framework for R. R package version 0.14.2, URL https://CRAN.R-project.org/package=shiny.

Cori A (2013). EpiEstim: Package to Estimate Time Varying Reproduction Numbers from Epidemic Curves. R package version 1.1-2, URL https://CRAN.R-project.org/package=EpiEstim.

Doi K, Sakabe K, Taruri M (2016). epifit: Flexible Modelling Functions for Epidemiological Data Analysis. R package version 0.0.6, URL https://CRAN.R-project.org/package=epifit.

Fienberg S, Wasserman S (1981). Analyzing Data from Multivariate Directed Graphs: An Application to Social Networks, pp. 289–306. John Wiley, London.

Frank O, Strauss D (1986). “Markov Graphs.” Journal of the American Statistical Association, 81, 832–842.

Garnett G, Anderson R (1993). “Factors Controlling the Spread of HIV in Heterosexual Communities in Developing Countries: Patterns of Mixing Between Different Age and Sexual Activity Classes.” Philos Trans R Soc Lond B Biol Sci, 342(1300), 137–59.

Garnett GP, Cousens S, Hallett TB, Steketee R, Walker N (2011). “Mathematical Models in the Evaluation of Health Programmes.” Lancet, 378(9790), 515–25.

Geyer C, Thompson E (1992). “Constrained Monte Carlo Maximum Likelihood for Dependent Data.” Journal of the Roal Statistical Society Series B, 54, 657–699.

Goodreau S, Carnegie N, Vittinghoff E, Lama J, Sanchez J, Grinsztejn B, Koblin B, Mayer K, Buchbinder S (2012). “What Drives the US and Peruvian HIV Epidemics in Men Who Have Sex With Men (MSM)?” PLoS ONE, 7(11), e50522.

Goodreau S, Cassels S, Kasprzyk D, Montano D, Greek A, Morris M (2010). “Concurrent Partnerships, Acute Infection and HIV Epidemic Dynamics Among Young Adults in Zimbabwe.” AIDS and Behavior, 16(2), 312–322.

Goodreau SM (2011). “A Decade of Modelling Research Yields Considerable Evidence for the Importance of Concurrency: A Response to Sawers and Stillwaggon.” Journal of the International AIDS Society, 14, 12.

Groendyke C, Welch D (2016). epinet: Epidemic/Network-Related Tools. R package version 2.1.7, URL https://CRAN.R-project.org/package=epinet.

Hallett T, Singh K, Smith J, White R, Abu-Raddad L, Garnett G (2008). “Understanding the Impact of Male Circumcision Interventions on the Spread of HIV in Southern Africa.” PLoS ONE, 3(5), e2212.

Handcock M, Hunter D, Butts C, Goodreau S, Morris M (2008). “statnet: Software Tools for the Representation, Visualization, Analysis and Simulation of Network Data.” Journal of Statistical Software, 24(1), 1548.

Handcock M, Robins G, Snijders T, Moody J, Besag J (2003). “Assessing degeneracy in statistical models of social networks.” Technical report, University of Washington Working Paper.

Handcock MS, Hunter DR, Butts CT, Goodreau SM, Krivitsky PN, Bender-deMoll S, Morris M (2015). statnet: Software Tools for the Statistical Analysis of Network Data. R package version 2016.9, URL CRAN.R-project.org/package=statnet.

Handcock MS, Hunter DR, Butts CT, Goodreau SM, Krivitsky PN, Morris M (2017). ergm: Fit, Simulate and Diagnose Exponential-Family Models for Networks. R package version 3.7.1, URL http://CRAN.R-project.org/package=ergm.

Hethcote HW, Van Ark JW (1987). “Epidemiological Models for Heterogeneous Populations: Proportionate Mixing, Parameter Estimation, and Immunization Programs.” Mathematical Biosciences, 84(1), 85–118.

Höhle M, Meyer S, Paul M (2016). surveillance: Temporal and Spatio-Temporal Modeling and Monitoring of Epidemic Phenomena. R package version 1.11.0, URL https://CRAN.R-project.org/package=surveillance.

Holland P, Leinhardt S (1981). “An Exponential Family of Probability Distributions for Directed Graphs.” JASA, 77, 33–50.

Hunter D, Handcock M (2006). “Inference in Curved Exponential Family Models for Networks.” Journal of Computational and Graphical Statistics, 15(3), 565–583.

Hunter D, Handcock M, Butts C, Goodreau S, Morris M (2008). “ergm: A Package to Fit, Simulate and Diagnose Exponential-Family Models for Networks.” Journal of Statistical Software, 24(3), 1–29.

Jenness S, Goodreau S, Rosenberg E, Beylerian E, Hoover K, Smith D, Sullivan P (2016a). “Impact of the Centers for Disease Control’s HIV Preexposure Prophylaxis Guidelines for Men Who Have Sex With Men in the United States.” Journal of Infectious Diseases, 214(12), 1800–1807.

Jenness SM, Goodreau SM, Morris M, Cassels S (2016b). “Effectiveness of Combination Packages for HIV-1 Prevention in Sub-Saharan Africa Depends on Partnership Network Structure.” Sexually Transmitted Infections, 92(8), 619–624.

Keeling MJ, Rohani P (2008). Modeling Infectious Diseases in Humans and Animals. Princeton University Press.

Khan B, Dombrowski K, Saad M (2014). “A Stochastic Agent-Based Model of Pathogen Propagation in Dynamic Multi-Relational Social Networks.” Simulation, 90(4), 460–484.

Koehly LM, Goodreau SM, Morris M (2004). “Exponential Family Models for Sampled and Census Network Data.” Sociological Methodology, 34(1), 241–270.

Kolaczyk ED, Krivitsky PN (2015). “On the Question of Effective Sample Size in Network Modeling: An Asymptotic Inquiry.” Statistical Science, 30(2), 184–198.

Krivitsky P, Handcock M (2014). “A Separable Model for Dynamic Networks.” Journal of the Royal Statistical Society B, 76(1), 29–46.

Krivitsky P, Handcock M, Morris M (2011). “Adjusting for Network Size and Composition Effects in Exponential-Family Random Graph Models.” Statistical Methodology, 8(4), 319–339.

Krivitsky P, Morris M (2017). “Inference for Social Network Models from Egocentrically-Sampled Data, with Application to Understanding Persistent Racial Disparities in HIV Prevalence in the US.” Annals of Applied Statistics, 11(1), 427–455.

Krivitsky PN, Handcock MS (2015). tergm: Fit, Simulate and Diagnose Models for Network Evolution Based on Exponential-Family Random Graph Models. R package version 3.4.0, URL http://CRAN.R-project.org/package=tergm.

Lessler J, Cummings DAT (2016). “Mechanistic Models of Infectious Disease and Their Impact on Public Health.” American Journal of Epidemiology, 183(5), 415–22.

Leung K, Kretzschmar M, Diekmann O (2015). “SI Infection on a Dynamic Partnership Network: Characterization of R0.” Journal of Mathematical Biology, 71(1), 1–56.

Morris M (1991). “A Log-Linear Modeling Framework for Selective Mixing.” Mathematical biosciences, 107(2), 349–77.

Morris M (1997). “Sexual Networks and HIV.” AIDS, 11 Suppl A, S209–16.

Morris M (2004). Network Epidemiology: A Handbook for Survey Design and Data Collection. Oxford University Press.

Morris M, Kretzschmar M (2000). “A Microsimulation Study of the Effect of Concurrent Partnerships on the Spread of HIV in Uganda.” Mathematical Population Studies, 8(2), 109–133.

Morris M, Kurth A, Hamilton D, Moody J, Wakefield S (2009). “Concurrent Partnerships and HIV Prevalence Disparities by Race: Linking Science and Public Health Practice.” American Journal of Public Health, 99(6), 1023–1031.

Poole D, Raftery AE (2000). “Inference for Deterministic Simulation Models: The Bayesian Melding Approach.” Journal of the American Statistical Association, 95(452), 1244–1255.

Santos Baquero O, Silveira Marques F (2015). EpiDynamics: Dynamic Models in Epidemiology. R package version 0.3.0, URL https://CRAN.R-project.org/package=EpiDynamics.

Schweinberger M (2011). “Instability, Sensitivity, and Degeneracy of Discrete Exponential Families.” Journal of the American Statistical Association, 106(496), 1361–1370.

Sullivan PS, Carballo-Dieguez A, Coates T, Goodreau SM, McGowan I, Sanders EJ, Smith A, Goswami P, Sanchez J (2012). “Successes and Challenges of HIV Prevention in Men Who Have Sex with Men.” Lancet, 380(9839), 388–99.

Toni T, Welch D, Strelkowa N, Ipsen A, Stumpf MPH (2009). “Approximate Bayesian Computation Scheme for Parameter Inference and Model Selection in Dynamical Systems.” Journal of the Royal Society Interface, 6(31), 187–202.

Wasserman S, Pattison P (1996). “Logit Models and Logistic Regressions for Social Networks: I. An Introduction to Markov Graphs and p*.” Psychometrika, 60, 401–426.

